# Why do some predicted protein structures fold poorly? Benchmarking AlphaFold, ESMFold, and Boltz in maize

**DOI:** 10.1101/2025.07.05.663230

**Authors:** Olivia C. Haley, Laura Tibbs-Cortes, Rita K. Hayford, Stephen Harding, Margaret Woodhouse, Ethalinda Cannon, Jack Gardiner, John Portwood, Taner Z. Sen, Hye-Seon Kim, Carson Andorf

**Affiliations:** Corn Insects and Crop Genetics Research Unit, US Department of Agriculture – Agricultural Research Service, Ames, IA 50011, USA; Division of Animal Sciences, University of Missouri, Columbia, MO 65211, US; Crop Improvement Genetics Research Unit, US Department of Agriculture – Agricultural Research Service, Albany, CA, 94710, USA; Department of Bioengineering, University of California, Berkeley, CA, 94720, USA; Mycotoxin Prevention and Applied Microbiology Research Unit, US Department of Agriculture – Agricultural Research Service, Peoria, Illinois 61604-3999

**Keywords:** Benchmarking, AlphaFold, Boltz, ESMFold, plant protein

## Abstract

Protein structure prediction tools have significantly reduced the time and cost to generate protein structures and accelerated protein discovery and design. However, plant proteins are underrepresented in sequence and structural datasets used to train these programs. To quantify the downstream impact of this deficiency, we benchmarked five structure-prediction programs (AlphaFold 2, AlphaFold 3, ESMFold, Boltz-1, and Boltz-2) across 417 well-characterized *Zea mays* genes. These “classical” genes represent a set of well-studied genes with known genetic and phenotypic effects. We generated structures for each gene using these programs and compared how sequence, structural, and evolutionary conservation impacted the structures’ confidence and geometric features. Proteins lacking conserved sequence and/or structural domains had on average 25% to 43% lower confidence scores than proteins having both domains. Proteome-wide phylostratigraphy revealed that species-specific proteins had substantially lower confidence scores than proteins conserved amongst angiosperms and Eukaryotes. Boltz-1 and ESMFold structures had the highest occurrence of structures with severe geometry issues, including overlapping atoms and unlikely bond angles. We also compared computational and experimental alignments of Arabidopsis, maize, rice, wheat, and soybean proteins from the Protein Data Bank and identified structures showing incongruences with experimental data. This study challenges the assertion that protein folding has been completely ‘solved’, and urges more investigation into benchmarking and standardized evaluation frameworks to improve model performance and assessment in agricultural crops.

## INTRODUCTION

In 2020, AlphaFold, developed by Google DeepMind, demonstrated the ability to predict 3D protein structures at near-experimental accuracy^1^, a feat that earned its creators the Nobel Prize in Chemistry. The remarkable performance of AlphaFold 2 in the 14th Critical Assessment of Protein Structure Prediction (CASP14) experiments marked a substantial advancement in the field^2^. Large-scale releases of predicted proteomes further showed the potential of this approach, with 58% of all residues being predicted with high confidence, 36% with very high confidence, and 98.5% of proteins containing at least one confident domain^3^. Additional recent advancements in structure prediction tools have expanded the methodological landscape: RoseTTAFold introduced a three-track neural network to integrate sequence, pairwise, and spatial information^4^; ESMFold^5^, and OmegaFold^6^ leveraged protein language models trained exclusively on sequence data; Chai-1^7^ incorporated multimodal inputs including structural and functional data; and Boltz-1^8^, Boltz-2^9^, and AlphaFold 3^10^ employed denoising diffusion models for iterative structural refinement. Boltz-2 is the most recently released protein folding program, and differs from the first iteration in its new affinity module and training data.

Despite these significant advancements in protein structure prediction, the definition of “high-quality prediction” has become an ongoing topic of discussion in structural biology, particularly regarding benchmarking standards in plant species. Protein structure prediction models are trained on protein structures, sequences, or a combination of the two. The Protein Data Bank (PDB), a database of experimental determined protein structures^11^, is commonly used to train protein structure prediction algorithms. However, there is uneven representation of clades across the tree of life in the PDB. For instance, fewer than 4% of the protein structures in the PDB are from the *Viridiplantae* (‘green plants’) clade that contains algal species and land plants. Such uneven sequence representations across the tree of life also negatively impact downstream inference for models incorporating sequence, requiring fine-tuning to improve performance^12^.

Furthermore, while these models are transforming the way scientists predict and use protein structures, they also reveal important limitations in their stereochemical accuracy (3D arrangement of atoms), which may lead to inaccuracies in the overall function of a protein. For example, McDonald et al.^13^ found that AlphaFold 2 had a higher performance in predicting Cɑ backbone structures than OmegaFold or RoseTTAFold, but struggled to predict mixed secondary structures (e.g., ɑ, β, ɑ/β, ɑ+β) and had varying degrees of success in the arrangement of the peptides’ dihedral angles. Additionally, Boltz-1 has exhibited a propensity for more steric clashes^8^ because, at the time of this manuscript, it does not incorporate any method to reduce the prevalence of clashes (unlike AlphaFold 3). Thus, given the underrepresentation of plant structures in structural databases such as the PDB, low annotation rates of plant sequences in curated databases, and the potential for problematic stereochemistry, assessing model performance on plant proteins is necessary to support protein structure prediction and engineering initiatives. Such assessments are essential for determining which methods/programs are a good fit for the data and offer the most promising structures to fulfill research goals.

Maize (*Zea mays* subsp. *mays*) presents a compelling case for further investigation into improving protein structure prediction models for crops given its critical role in global food production^14^ and a substantial array of genomic sources, including comprehensive pangenomic data^15^, detailed mappings of stress response networks^16^, extensive datasets of natural genomic variation^17^, and vast amount of gene expression data^18^. As a model organism^19^, maize has been heavily studied to provide genomic insights into domestication, organogenesis, C4 photosynthesis, and cellular fate determination, amongst other biological processes^20^. Significant investments and technological advancements in sequencing and breeding technologies have further accelerated maize genetic research and breeding effort ^21^. However, crops like maize exhibit some of the lowest prediction confidences in the AlphaFold database^22^. Once more, benchmarking and critical assessments of the performance of structure prediction programs are still sparse for plant-specific proteins^23^.

While protein structure prediction tools are accessible through online platforms, their use is often limited by sequence length, the number of sequences allowed per session, time limits, and memory constraints. Due to these limitations, conducting a comprehensive, proteome-wide structural analysis is not feasible without access to GPU clusters. To address this, we present a case study assessing the performance of five protein structure prediction programs (AlphaFold 2, AlphaFold 3, ESMFold, Boltz-1, and Boltz-2) for 417 “classical” genes in maize^24^ defined as having well-annotated functions and phenotypes^20^. This subset of structure prediction programs was selected because of their prominence in the literature (AlphaFold 2, ESMFold) and novelty (AlphaFold 3, Boltz-1, Boltz-2). During this exploration, we quantified several aspects of the predicted protein structures. First, we assessed the quality of the unrefined structures generated by each method from a stereochemical perspective, including evaluations of geometry, bond angles, and other structural features. We also examined how structural characteristics, sequence properties, and evolutionary conservation influenced each method’s confidence scores. Finally, we compared both global and local structural alignments between selected experimentally determined plant protein structures and their corresponding in silico predictions. This assessment highlights that confident protein structure predictions remain challenging for young and maize-specific proteins, as well as those lacking conserved sequence or structural domains, across both maize and other model species. It also demonstrates that different prediction programs offer distinct advantages: ESMFold is the fastest, AlphaFold produces the most confident predictions, and the Boltz programs provide a good compromise, guiding their strategic deployment within bioinformatics pipelines.

## MATERIALS & METHODS

### Dataset retrieval

The protein sequence corresponding to the canonical transcript for each maize classical gene was downloaded from MaizeGDB^25,26^. The sequences are included in the **Supplemental Data**. The experimentally resolved plant proteins were retrieved by querying the PDB^11^ based on the taxonomy identifier for: *Arabidopsis thaliana* (Arabidopsis, Taxon ID: 3702); *Oryza sativa* (rice, Taxon ID: 4530); *Triticum aestivum* (bread wheat, Taxon ID: 4565); *Zea mays* (maize, Taxon ID: 4577); *Glycine max* (soybean, Taxon ID: 3847). Only experimental structures with an R_free_ value ≤ 0.30 and sequence similarity ≥ 95% to the reference protein sequence in UniProt were used in the analysis.

### Calculating the pLDDT of model organism structures

The AlphaFold Protein Structure Database^1,22^ provides folded protein structures for the proteomes of 16 model organisms across the kingdoms of life: Archaebacteria (*Methanocaldococcus jannaschii*), Eubacteria (*Escherichia coli*), Protista (*Dictyostelium discoideum*), Fungi (*Candida albicans*, *Saccharomyces cerevisiae*, *Schizosaccharomyces pombe*), Plantae (*Arabidopsis thaliana*, *Glycine max*, *Oryza sativa*, *Zea mays*), and Animalia (*Homo sapiens*, *Danio rerio*, *Mus musculus*, *Rattus norvegicus*, *Drosophila melanogaster*, *Caenorhabditis elegans*).

The AlphaFold 2 structures from 16 model organisms were downloaded from the AlphaFold Protein Structure Database (version 4), and the per-residue pLDDT scores were extracted using custom scripts in Python (version 3.11). The means of the per-residue pLDDT scores were calculated in Python to generate the structure-averaged pLDDT scores.

### Graphics Processing Units (GPUs) and Runtime Evaluation

The GPU computing environment for this study was provided by the Atlas High Performance Computing Collaboratory hosted by Mississippi State University, through agreements with SCINet Scientific Computing Office. Atlas is a Cray CS500 Linux cluster which, at the time of writing this manuscript, has 11,520 2.40GHz Xeon Platinum 8260 processor cores, 101 terabytes of RAM, 8 NVIDIA V100 GPUs, 5 HPE Apollo XL675d nodes equipped with 8 NVIDIA A100 GPUs per node, 12 NVIDIA L40S GPU nodes, each with 4 GPUs with 48 GB of VRAM and 91.6 TeraFLOPS, and a Mellanox HDR100 InfiniBand interconnect. The peak performance of Atlas is currently 565 TeraFLOPS.

To assess factors influencing program runtimes, protein structure predictions were performed individually on a single GPU node, on either a V100 node (32 GB per GPU), A100 node (80 GB per GPU), or an L40S node (48 GB per GPU). The time taken to generate each protein structure was recorded using the SLURM workload manager.

The structures generated on the A100s were used for further analysis of the methods’ protein folding performance (i.e., steric hindrances, structural alignments). AlphaFold 2 was exclusively run on A100 nodes because its database is stored on local solid-state drives, meaning structures from the V100 or L40S nodes are not included in structural comparisons or further analyses. All the structures are still available in the GitHub repository.

### CATH Domain Analysis

CATH is a hierarchical classification system for protein structures based on structural domains from the PDB. The CATH hierarchy is organized into four levels: Class (C), which corresponds to the secondary structure content; Architecture (A), the three-dimensional secondary structure arrangement; Topology (T), which integrates information into the arrangement of the secondary structure elements; and Homologous superfamily (H) domain relation based on evolutionary information.

CATH domain analysis was used to determine the influence of structural conservation on the programs’ runtime, and subsequent confidence in the generated protein structure. The CATH domain(s) corresponding to each classical gene were retrieved from the online database using the API service^27^. The database was queried using the proteins’ sequence as input to identify conserved structural domains and functional families. The Python IntervalTree package was used to identify non-overlapping domains based on the significance values provided during the query.

### Pfam Domain Analysis

Akin to CATH domains, Pfam domains are widely used in computational biology, and consist of highly conserved sequences that have often been manually annotated with information regarding the domain’s function^28^. The presence or absence of protein family (Pfam) domains was used to investigate the effect of sequence conservation on the runtime of the programs and on structure confidence. The Pfam annotations of the classical genes were downloaded from MaizeGDB.

### Evolutionary Conservation Analysis

In evolutionary biology, phylostratigraphy is a technique used to estimate the age of genes by following their origins via a phylogenetic tree and allocating each gene to a “phylostratum” according to the most ancient common ancestor that has a homolog^29^. The evolutionary ages of the gene models were predicted using Phylostratr, an R-based framework for estimating the evolutionary ages of gene models across an entire species^30^. The program features a fully automated analysis to infer phylostrata and diagnostics for quality control. In Phylostratr, a genes’ evolutionary origin is estimated by searching for homologs within representative proteomes from increasingly broad clades. The ‘youngest’ clade that contains a homolog of the gene is that gene’s phylostratum.

The phylostrata were calculated for B73 canonical proteins using the R package Phylostratr. Proteomes were downloaded from UniProt in December 2024. The eukaryotic and prokaryotic trees were created using modified versions of the ‘diverse_subtree’ and ‘uniprot_sample_prokaryotes’ functions, respectively. The Phylostratr scores were used to assess the impact of evolutionary conservation on the model’s pLDDT scores.

### Global and Local Structural Alignments

The FoldSeek command line program^31^ was used to perform the structural alignments between the *in silico* predicted structures and the experimentally-resolved structures. To perform the alignments, the PDB structure files were used to form a ‘ground truth’ reference database which was then queried using the computational structures and default parameters: -s 9.5; --num-iterations 0; --max-seqs 1000; -e 0.001; --alignment-type 2 (3Di+AA); -c 0.0; --cov-mode 0.

### Assessment of Steric Hindrances

The online SWISS-MODEL Structure Assessment Tool^32^ was used to perform the stereochemistry evaluations. This program was chosen because it simultaneously calculates and visualizes multiple stereochemical features using the MolProbity program^33^, Ramachandran plots^34^, secondary structure content using DSSP^35^, statistical model quality assessment with QMEANDisCo^36^, and predicts if the protein is likely localized to the cellular membrane.

### Data Analyses

We first assessed computational performance by modelling structure-generation time as a function of protein length and extrapolating those regressions to estimate how long each program would require to fold an entire proteome. We then examined model confidence. Two-sample t-tests compared predictions for proteins that contain conserved sequence or structural domains with those that do not, while a one-way ANOVA evaluated whether high-confidence predictions persist when a conserved domain is missing altogether. To relate confidence to evolutionary conservation at the gene level we calculated Spearman rank correlations, and to determine whether confidence varies across discrete conservation strata we applied a Kruskal– Wallis test followed by Dunn’s post-hoc comparisons. Stereochemical accuracy was analyzed next: linear mixed-effects models tested whether the tools generate structures with comparable steric architectures, and a principal-component analysis was performed to uncover broader trends in those steric features across programs. Finally, overall structural accuracy was benchmarked by fitting an additional linear mixed-effects model that compared predicted structures with experimentally determined plant protein structures deposited in the Protein Data Bank.

## RESULTS & DISCUSSION

### AlphaFold Database reference proteomes reveal lower performance for grass species

The Protein Data Bank (PDB) is a database of protein structures which were resolved using experimental methods such as X-ray crystallography or nuclear magnetic resonance (NMR). The structures in this database are often used to train programs used for structure prediction and protein design. However, plant species are significantly underrepresented in the PDB. Proteins from the Viridiplantae (‘green plants’) clade constitute only 3.40% (8,017/ 236,060) of experimental structures in the PDB as of May 2025. In contrast, human protein structures make up 31.5% (74,433/236,060) of the PDB, despite having a substantially smaller proteome (∼20,000 proteins) compared to species like maize or wheat (∼40,000 and ∼80,000 proteins, respectively).

The consequences of this bias are apparent in the AlphaFold Structure Database, particularly for maize and rice, where 43% to 51% of their proteins have an average confidence score below the high-confidence threshold (pLDDT score < 70; **Figure 1**). In humans, only 32.8% of the AlphaFold 2 structures have pLDDT scores in this range. However, the disproportionate representation of plant proteins in reference databases does not appear to affect all plant species. Only 29.9% of the AlphaFold 2 Arabidopsis structures had structure-averaged pLDDT scores below the high-confidence threshold. This distribution is not surprising since there are almost 10 times more experimental Arabidopsis structures (n = 2,395) in the PDB than maize (n = 272) or rice (n = 347). Nevertheless, soybean, which has even fewer experimental structures (n = 187), had a greater proportion of high-confidence residues and structures than the grasses. We hypothesize that this observation is due to the closer phylogenetic relationship between soybean and Arabidopsis, both of which are dicot species. Given this evolutionary proximity, proteins from Arabidopsis are more likely to share conserved sequence and structural features with those of soybean, thereby providing broader and more accurate structural coverage. In contrast, maize and rice are monocot species from a more distantly related clade, thus their proteins may exhibit structural divergences that are underrepresented in the Arabidopsis-derived entries in the Protein Data Bank (PDB). These evolutionary differences likely contribute to the reduced applicability of Arabidopsis-based structural models to monocot proteins.

**Figure 1.**
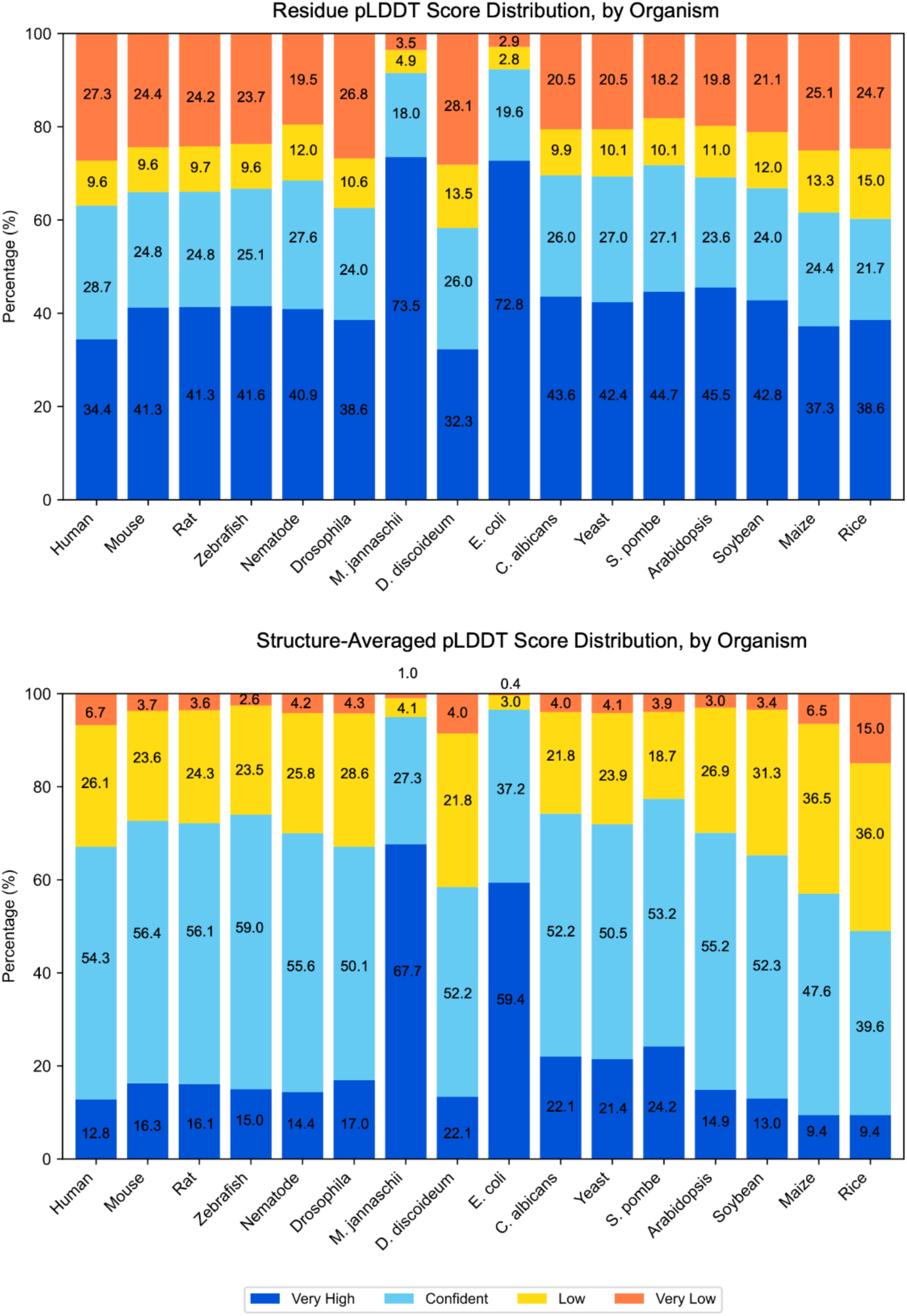
The distribution of AlphaFold 2 confidence scores by residue (***top***), and averaged across the proteins’ structure (***bottom***) for 16 model species in the AlphaFold Structure Database.

This analysis also does not account for proteins with intrinsic disorder, which can also result in low per-residue pLDDT scores^1^. Yet, these findings substantiate the need to further investigate factors that affect the programs’ ability to fold proteins whose structures and sequences were not seen during training.

### Benchmarking Runtime and Memory Demands of Protein Structure Prediction

Understanding a program’s runtime and memory requirements is important to supporting research initiatives in computational biology. In this field, research workflows often involve large, complex datasets, whose analyses can become time- and resource-consuming if not optimized. Computational resources, particularly specialized hardware like GPUs, may also have limited availability or incur significant costs, adding complexity to project planning and resource allocation. Ultimately, runtime is not just a technical detail, it’s a critical component of methodological selection. By evaluating the trade-offs between the runtime and downstream accuracy of the generated structure model, we aim to assist researchers in the protein structure space to make more informed decisions about program selection.

The program runtimes on NVIDIA’s L40S, A100 and V100 GPU cores are summarized in **Figure 2**. Because the runtimes were significantly shorter for ESMFold,Boltz-1, and Boltz-2 compared to AlphaFold 2 and AlphaFold 3, direct comparisons were made between ESMFold, Boltz-1, and Boltz-2, which were separate from AlphaFold 2 and AlphaFold 3. A tabulated summary is available in **Supplemental Table 1**, and the full dataset is available as **Supplemental Dataset**. Occasionally, the programs would fail to produce a predicted structure, which was usually due to out-of-memory errors caused by a large number of sequence hits that would increase the memory required for the multiple-sequence alignment (MSA) construction stage. For example, Zm00001eb376160 had the largest number (12,123) of sequence hits from the Big Fantastic Database (BFD), and its structure was only generated by Boltz-1, Boltz-2, and ESMFold. Memory issues would also occur when generating AlphaFold 2 structures for longer proteins (>1,000 residues). These out-of-memory instances would not be included in the runtime or subsequent analyses.

**Figure 2.**
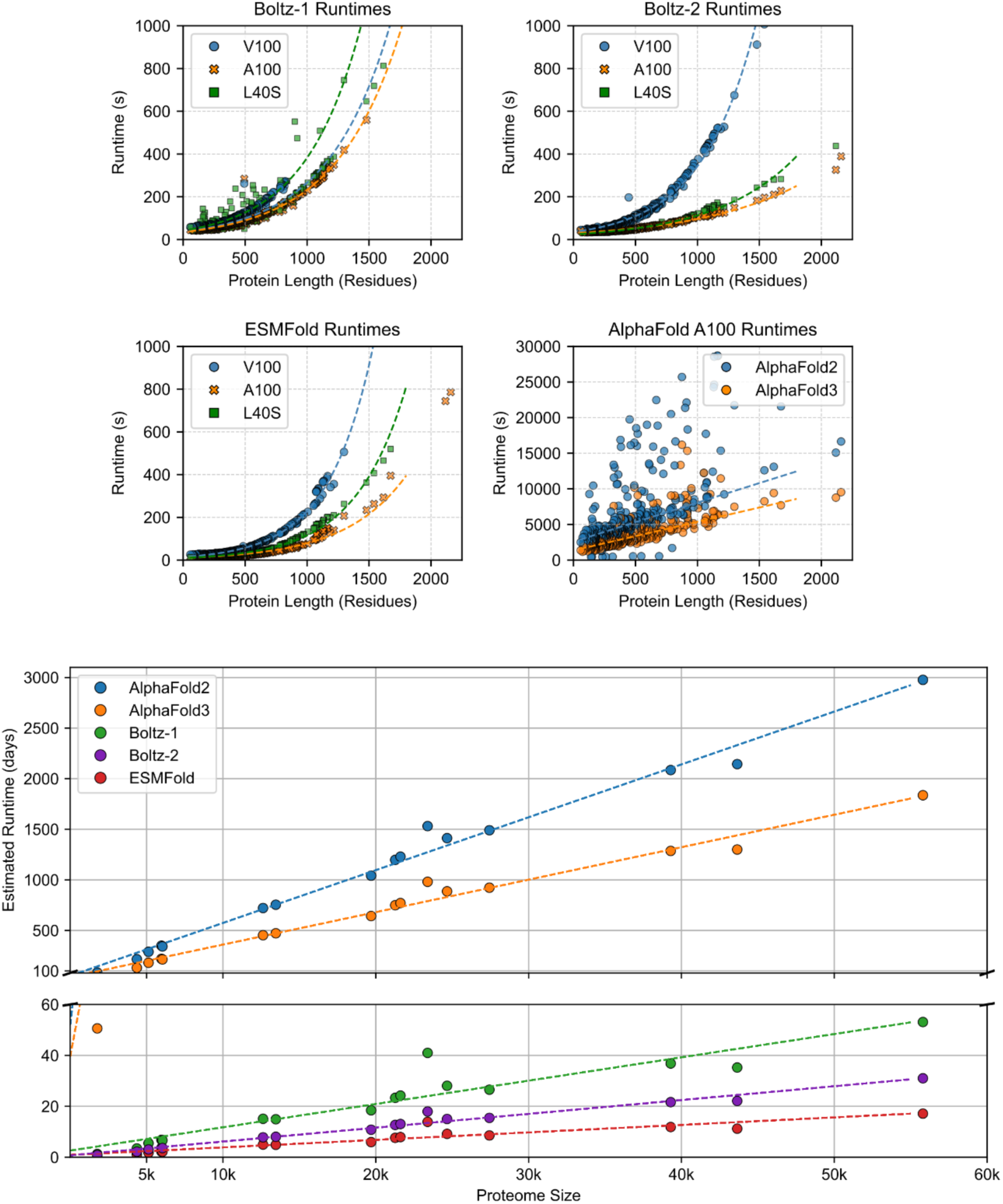
Runtimes for the structure prediction programs were benchmarked on A100, V100, and L40S GPUs (top). All programs were benchmarked on the A100 GPUs, while only ESMFold,Boltz-1, and Boltz-2 were benchmarked using the V100 and L40S GPUs (top). Linear or exponential regression models were used to model the relationship between protein length and runtime, and to estimate total runtime for 16 representative proteomes on a single A100 GPU (bottom). Note the broken axis, and that the estimated runtimes for the proteomes are in days.

### Runtime Benchmarking: ESMFold produces the fastest predictions

ESMFold consistently generated the protein structures faster than either Boltz program on both the A100 and V100 GPUs, and both models generated the structures substantially faster than AlphaFold 2 or AlphaFold 3. On the A100 cores, for example, ESMFold generated the protein structures ∼170 times faster than AlphaFold 2 and ∼3 times faster than Boltz-1. The speed of ESMFold is expected, as it is based on a language model; ESMFold does not require a multiple sequence alignment (MSA) against large sequence databases and thus avoids this resource-intensive bottleneck. The ‘optimized’ MSA generation steps of the Boltz-1 and AlphaFold 3 models also appear to have reduced the runtime required to generate the structures as compared to AlphaFold 2. AlphaFold 3 also generated more of the structures than AlphaFold 2 with a total inference time 1.3 times faster than AlphaFold 2.

It is well-documented that protein length is the primary explanatory factor for how long the programs take to generate a protein structure^1,5^. Unsurprisingly, protein length was strongly correlated with structure generation time, as assessed using both linear (Pearson *r*) and non-parametric (Spearman rank) correlation analyses. Predictive models were developed to estimate the time required to fold an entire proteome based on proteome size (**Figure 3**). To fit the data, the data were first split 80/20 into training and validation sets, with the training set used to estimate model parameters and the validation set used to assess model performance. Based on initial visualizations, linear relationships were assumed for AlphaFold 2 and AlphaFold 3, while exponential relationships were explored for Boltz-1, Boltz-2, and ESMFold. Due to the greater variability (‘noise’) in AlphaFold 2 and AlphaFold 3 predictions, outliers were removed using the interquartile range method. Briefly, the IQR is the range between the first quartile (Q1, 25th percentile) and third quartile (Q3, 75th percentile). Any data point below Q1 − 1.5×IQR or above Q3 + 1.5×IQR is considered an outlier and removed from the dataset to reduce noise during fitting.

**Figure 3.**
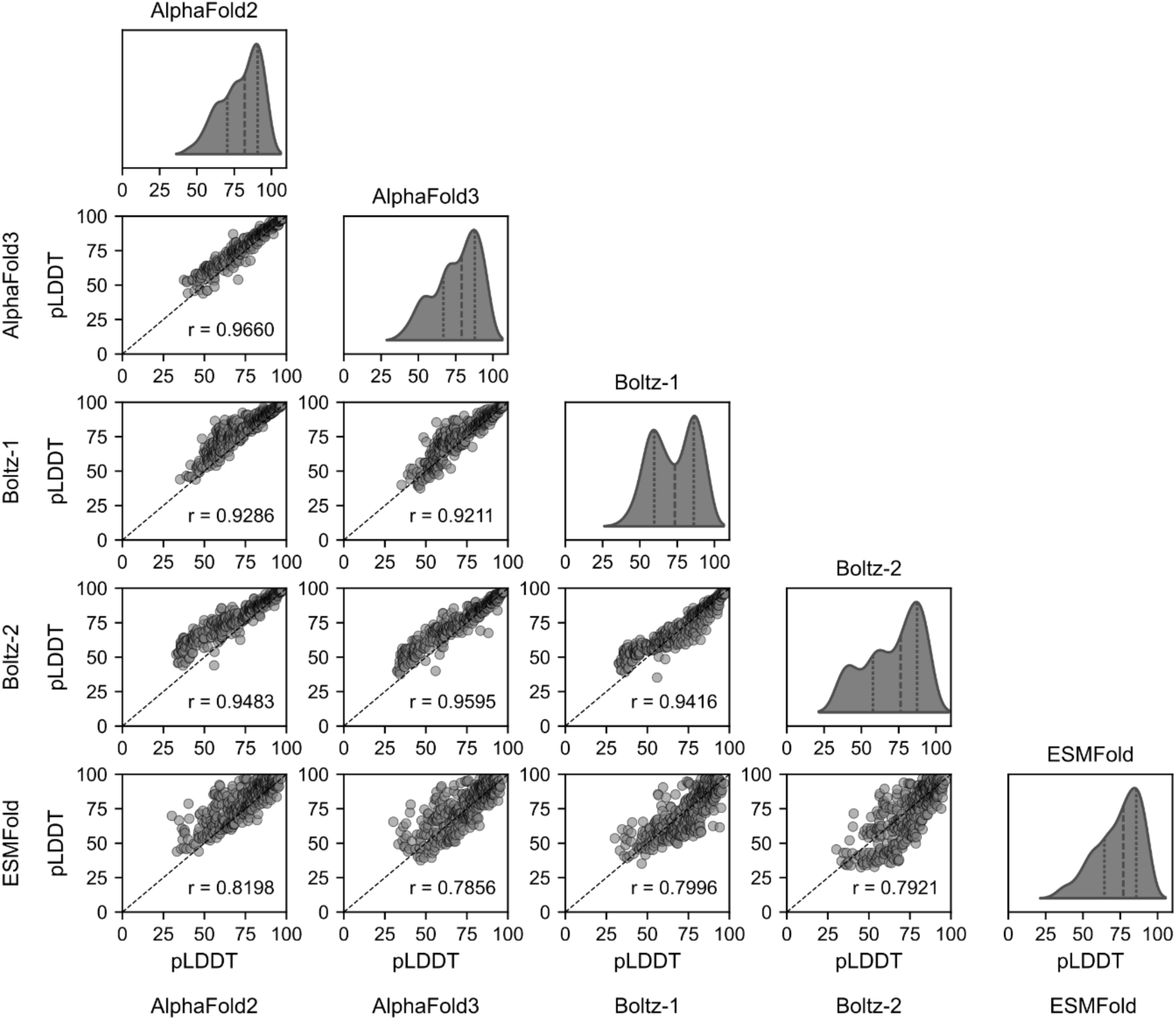
To visualize the distribution of the models’ confidence, the per-residue pLDDT scores were averaged for each structure generated by AlphaFold 2, AlphaFold 3, Boltz-1, Boltz-2, and ESMFold. The distribution of the pLDDT scores indicates that AlphaFold 2 was generally more confident in its generated structures than any other program. The Pearson correlation coefficient (*r*) for pairwise comparisons is also shown for each comparison.

The regression analyses indicated that runtime for ESMFold, Boltz-1, and Boltz-2 was largely explained by protein length on all three GPU cores (R^2^: 0.8172 to 0.9926); **Table 1**). In contrast, the AlphaFold 2 and AlphaFold 3 programs experienced more variation that required outlier reduction. Still, for AlphaFold 2, the R^2^ value (0.5378) indicated that the variation was not well explained by protein length alone. Thus, there may be a more complex relationship between the proteins’ characteristics and time required to generate a 3D structure for AlphaFold 2 and AlphaFold 3.

**Table 1.** A tabulated summary of the linear regression metrics for the predictive models developed to describe the relationship between the proteins’ length (amino acid residues) and the time required to generate its structure (s) for each of the programs and GPUs.

| Program<sup>a</sup> | GPU<sup>b</sup> | Fit<sup>c</sup> | RMSE<sup>d</sup> | R<sup>2</sup><sup>e</sup> |
| --- | --- | --- | --- | --- |
| AlphaFold 2 | A100 | Linear | 119.3 s | 0.5378 |
| AlphaFold 3 | A100 | Linear | 463.5 s | 0.8335 |
| Boltz-1 | A100 | Exponential | 3.7 s | 0.9926 |
|  | V100 | Exponential | 21.7 s | 0.9765 |
|  | L40S | Exponential | 22.4 s | 0.8172 |
| Boltz-2 | A100 | Exponential | 22.4 s | 0.9252 |
|  | V100 | Exponential | 16.8 s | 0.8893 |
|  | L40S | Exponential | 31.0 s | 0.9473 |
| ESMFold | A100 | Exponential | 11.2 s | 0.9494 |
|  | V100 | Exponential | 9.89 s | 0.9857 |
|  | L40S | Exponential | 11.5 s | 0.9708 |

<sup>a</sup> The structure prediction program
<sup>b</sup> The type of graphics processing unit
<sup>c</sup> The regression model used to fit the curve
<sup>d</sup> The root mean squared error of the model measured in seconds
<sup>e</sup> The coefficient of determination of the model: proportion of variance in the response variable that is explained by the fitted model (ranges from 0 = no explanatory power to 1 = perfect fit)

### Model Confidence: The AlphaFold methods produce the highest confidence models

Despite its speed, ESMFold also had the lowest confidence in its structures, according to the distribution of the confidence (predicted Local Distance Difference Test; pLDDT) scores (**Figure 3**). The metric was developed by the makers of AlphaFold to assess how confident the program was in its generated structure, by comparing the structure to a template protein (often from the PDB). It has since become a common initial assessment of structure quality, though various studies report it may overestimate or underestimate model quality^37^ or be skewed by intrinsically disordered regions^3^. AlphaFold 2, AlphaFold 3, Boltz-2, and Boltz-1 (in order of their overall confidence scores) generally produced more confident structures than ESMFold.

Additionally, the Pearson correlation coefficient was calculated between the pLDDT scores of the models to assess the harmonicity of their predictions. This analysis aimed to determine whether the pLDDT scores of one model tended to increase when the pLDDT scores of the other model increased; a high positive correlation could suggest that both models produce similar confidence in their predictions across the entire dataset. Conversely, a weak or negative correlation could indicate that the models’ pLDDT scores are not harmonic, meaning they do not consistently increase or decrease together. In this case, the models might diverge in their structural confidence assessments, potentially reflecting differences in their underlying prediction mechanisms. The Pearson correlation coefficient (*r*) for pairwise comparisons indicated a high linear correlation (0.7856 – 0.9660) between the methods’ pLDDT scores. Accordingly, when one model is more confident in its predictions, the other models also tend to be more confident in their predictions for the gene model. Interestingly, the pairwise pLDDT comparisons between Boltz-1 and Boltz-2 revealed a sigmoidal pattern, appearing as if the new features added to the Boltz program improved its performance for some proteins while reducing the performance for others. However, this is a point that must be explored using a larger, more comprehensive set of protein structures.

### Less conserved proteins were predicted with less confidence

From the distribution of the model confidence scores (**Figure 3**), it is evident that there were instances where the methods failed to generate highly confident structures. Thus, the factors driving the pLDDT scores were also explored, with an emphasis on testing the hypothesis that the scores would be lower for gene models that were not well-represented in 1) sequence-based databases, 2) structural databases, and/or 3) for gene models that were not evolutionarily conserved. These factors differ in that a gene model may be considered evolutionarily ‘young’ (i.e., arose in a recent genomic event such as gene duplication), but could still be well-represented in protein structure training data if its structure (or a homolog) was experimentally resolved.

The relationship between the models’ pLDDT scores and sequence/structure conservation was evaluated based on the presence/absence of either sequence-based (Protein Families; Pfam) and/or structure-based (CATH) domains. As previously mentioned, CATH is a hierarchical system that classifies protein structures based on: Class (secondary structure content), Architecture (3D arrangement), Topology (connectivity of structural elements), and Homologous superfamily (evolutionary relationships)^38^. Similarly, Pfam domains, which are widely used in computational biology, are highly conserved sequences, often manually annotated with functional information^28^.

Preliminary independent two-sample *t*-tests were conducted to assess whether there were statistically significant differences in the predicted structures’ confidence (pLDDT) scores between proteins with and without defined CATH or Pfam domains. As expected, regardless of the program/method, proteins with conserved CATH or Pfam domains consistently had significantly higher confidence scores than proteins lacking either (**Figure 4**). On average, structures containing CATH and Pfam domains had pLDDT scores that were higher by 21.2 units in AlphaFold 2, 19.5 units in AlphaFold 3, 21.2 units in Boltz-1, 24.1 units in Boltz-2 and 33.2 units in ESMFold, compared to structures without both domains. Post-hoc statistical tests were performed to see if there were different consequences of not having either CATH or Pfam domain, or if they were equally important to ensuring a high overall confidence score (**Supplemental Figure 1**).

**Figure 4.**
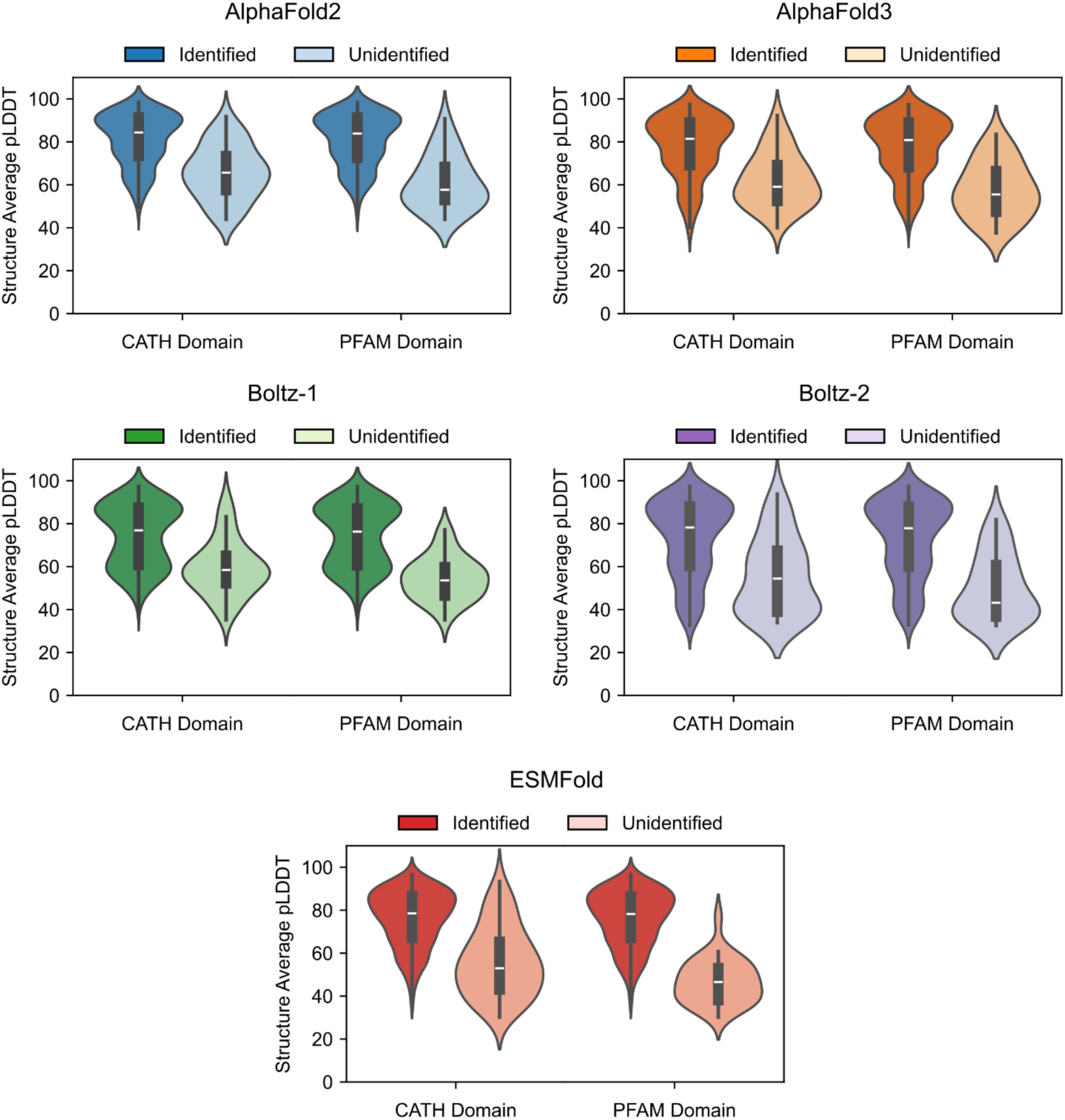
An analysis of methods’ confidence (pLDDT) scores based on the sequence/structural conservation of the protein and the evolutionary conservation of the gene model comparing when either a CATH or Pfam domain was identified (light color) vs unidentified (dark color). The presence/absence of the CATH and Pfam domains was used as an indicator of structural and sequence conservation, respectively. Within each program, proteins lacking either domain had statistically lower pLDDT scores (p < 0.05) than those having an identifiable domain.

To evaluate these individual and combined effects of domain presence, a one-way ANOVA was performed by grouping the proteins into four categories: (1) both domains present, (2) Pfam absent and CATH present, (3) Pfam present and CATH absent, and (4) both domains absent. The group variable was treated as a fixed effect. Post-hoc pairwise comparisons were conducted using the Bonferroni correction for multiple comparisons, with an adjusted significance threshold of α = 0.0083 (0.05 / 6).

The methods followed a similar pattern: proteins without either domain would have significantly lower confidence scores, but these scores would be statistically indistinguishable from proteins that were missing both domain types. For example, at 95% confidence, AlphaFold 2 structures containing both CATH and Pfam domains had pLDDT scores 5.3 to 32.1 units higher than structures with only a CATH domain (p < 0.0001), and 4.0 to 19.2 higher than those with only a Pfam domain (p < 0.001). However, there were no statistically significant differences in the pLDDT scores between the structures having only a CATH domain or only a Pfam domain (p=0.4447). In practice, this means that generating a structure for a plant protein with low sequence conservation is just as difficult as generating a structure for a protein with low structural conservation.

### All methods had lower confidence for evolutionarily young and species-specific proteins

In this study, model confidence decreased with evolutionary age (**Figure 5**), and the models struggled to produce highly confident structures for proteins encoded by gene models that were less conserved, with species-specific orphan proteins^39^ exhibiting the lowest confidence scores. Specifically, a Spearman correlation test between the Phylostratr scores and pLDDT scores was used to assess the overall relationship between evolutionary conservation and confidence scores. The correlation test was performed separately for each program. The correlation coefficients ranged between -0.5078 to -0.5315, indicating a moderate negative relationship between the evolutionary ages of the gene models and the confidence scores. However, the small sample size of gene models in the younger phylostrata limited the strength of this inference (there were only seven genes conserved at the *Poaceae/*grasses level, for example). To increase the power and generalizability of these analyses, we compared the pLDDT scores of AlphaFold 2 and ESMFold structures across the entire B73 proteome. These structures had previously been generated and are available at MaizeGDB^48^.

**Figure 5.**
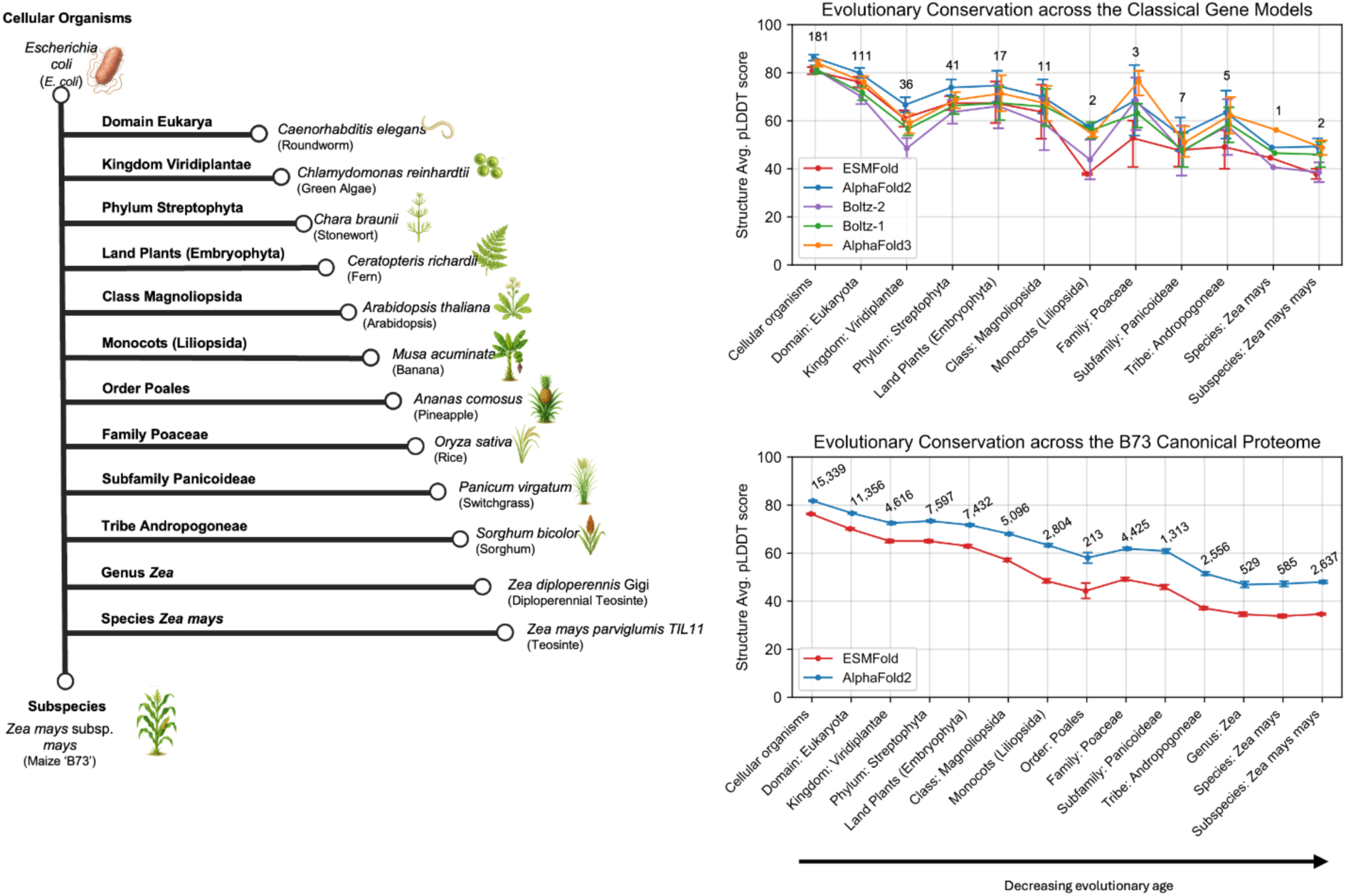
The structure-averaged pLDDT scores of the protein structures by phylostratigraphic age. The leftmost panel depicts the evolutionary relationship of the phylostrata, including a representative species at each phylostratigraphic level. The rightmost top panel shows the mean pLDDT scores for the protein structures generated from the classical genes, by each of the four methods. The lower panel shows the mean pLDDT score for each phylostratum for the in silico structures predicted by ESMFold and AlphaFold 2 for the entire B73 proteome. Error bars represent the 95% confidence intervals. Note that there were no classical gene models in the Order Poales, or Genus Zea, thus these strata do not appear on the x-axis in the rightmost top panel.

The pLDDT scores at each phylostratigraphic level were compared first within each program, and then across the programs. Within-program variation of pLDDT scores were measured using a Kruskal–Wallis nonparametric test to account for non-normal distributions and unequal group sizes; significant differences were further examined using Dunn’s post hoc test with Bonferroni adjustment (**Figure 5**). For both AlphaFold 2 and ESMFold, the pLDDT scores progressively decreased, with each level having statistically lower pLDDT scores than the last. There was one exception between the order *Poales* and *Poaceae* family where the pLDDT scores increased. This increase likely reflects the number of high-quality proteomes in this family such as rice, wheat, and maize. Between-program comparisons were performed using independent t-tests at each stratum and found that AlphaFold 2 consistently had higher confidence scores than ESMFold.

Many of the maize-specific proteins had low (<70) confidence scores (**Figure 5**). The average pLDDT score at this level was 48.0 for AlphaFold 2 structures, and 34.6 for ESMFold structures. The lower scores are logical, since these young genes are generally fast-evolving^40^ and are thus more likely to have higher sequence variation and lack conserved domains in structural databases. However, they are also highly valuable to future crop breeding efforts due to their involvement in tolerance to adverse conditions such as host adaptation to disease resistance^16^. To further enhance the utility of these protein structure predictions in agronomic research, it is essential to resolve the structures of these younger genes and integrate them into protein structure prediction methods.

### The methods generate structures with distinct steric features

It is common practice in advanced protein design studies to filter generated protein structures for steric clashes and stereochemical issues, by using tools such as Posebusters^41^, or to refine them with molecular dynamics simulations. However, some studies on computational protein structures forgo these steps due to the additional time and domain-specific expertise required. Therefore, it is essential for researchers to critically evaluate the stereochemistry and steric hindrances present in the methods’ raw, unrefined protein structures. We used the Molprobity module of the SWISS-Model Structural Assessment web tool to evaluate how well the predicted models obey the physical laws of protein structure and folding.

The MolProbity program provides multiple metrics, including the MolProbity score, clash score, rotamer outliers, Cβ deviations, Ramachandran favored residues, and Ramachandran unfavored residues. The ideal ranges for the metrics were determined based on recommendations from the literature, and are discussed below. The MolProbity score is a composite metric that combines the clash score, Ramachandran outliers, and rotamer outliers into a single score. The lower the MolProbity score, the better the model is expected to adhere to known chemical and physical constraints with < 2.0 being a cut-off previously used to denote overall good structural quality^33^. Clash score is the number of atomic overlaps between non-bonded atoms per 1000 atoms in the structure; high clash scores (>30) are suggestive of poor model geometry^42^. Cβ deviations are another indication of distorted backbone geometry, with the optimal number being 0^43^. Unfavorable side chain angles are indicated by: rotamer outliers (>0.5% indicates the presence of rare or physically unfavorable side chain conformations); Ramachandran favored (>98% is desired^43^); and Ramachandran outliers (>2% indicating that the structure may need review^42^).

Statistical differences in the stereochemical features of structures generated by the methods were assessed using a linear mixed-effects model, with the method as a fixed effect and classical gene as a random effect. Independent *t*-tests were then performed to quantify the differences between the individual models. As an overall indicator of quality, the MolProbity scores indicated AlphaFold 2 and AlphaFold 3 generally had fewer steric hindrances than ESMFold and Boltz-1 (**Figure 6**). Despite slightly lower confidence scores, AlphaFold 3 had fewer rotamer outliers and Cβ deviations (p < 0.05) than AlphaFold 2. Boltz-1 had the fewest rotamer outliers and lowest prevalence of Ramachandran outliers compared to the other models (p < 0.05), and its generated structures had fewer Cβ deviations than AlphaFold 2 or AlphaFold 3 (p < 0.05). However, the clash scores for Boltz-1’s structures were significantly (p < 0.05) higher than the other programs, including Boltz-2. The lower Clash scores and MolProbity scores for Boltz-2 compared to Boltz-1 indicated that the authors’ improvements to the program resulted in substantial improvements to the generated protein structures that may make them more suitable for downstream structure-based inference. In fact, the MolProbity scores between Boltz-2 and AlphaFold 3 were statistically indistinguishable, indicating that the performance of Boltz-2 is more comparable to AlphaFold 3 than Boltz-1. However, the MolProbity score also indicates that the quality of raw AlphaFold 2 structures is still higher than that of raw Boltz-2 structures.

**Figure 6.**
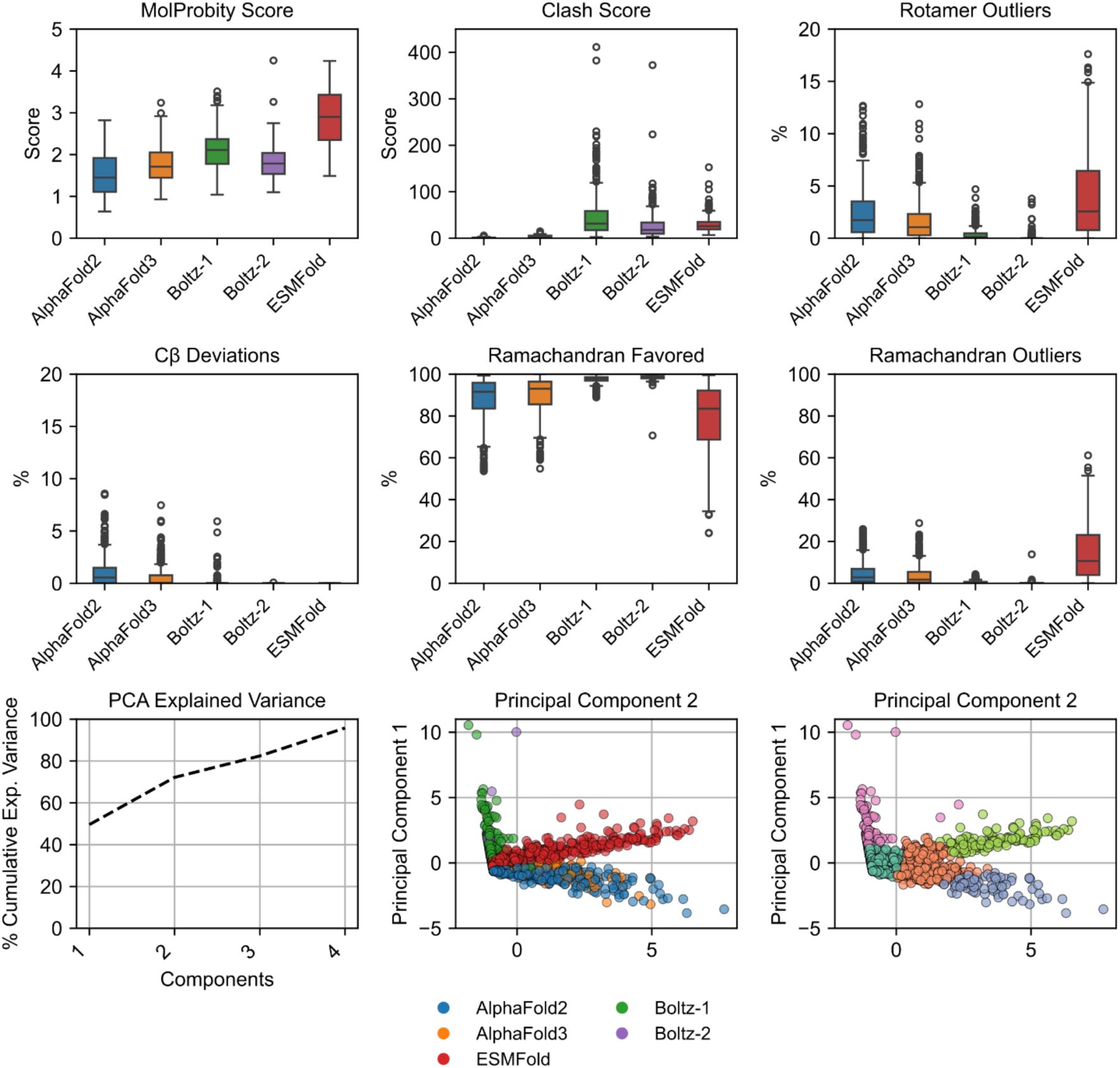
The distribution of a subset of the steric metrics, by the structure prediction program. The SWISSModel Expasy Structure Assessment web service was used to evaluate the stereochemistry of the generated structures. The metrics (excluding MolProbity score) were used as features in a Principal Component Analysis to test if the methods generated structures with distinct steric features. The explained variance ratio of the principal components, unclustered, and KMeans-clustered PCA coordinates are shown.

A Principal Component Analysis (PCA) was then performed using the structure scores to explore whether they can be used to distinguish between the structure prediction methods. The percentage of Ramachandran Favored residues were excluded from the feature matrix since its variance inflation factor (VIF) indicated a high degree of multicollinearity (VIF = 20.460378). While the MolProbity Score only had a moder degree of multicollinearity with the other features (4.699601), it was also removed from the dataset as its calculation involves using the clash score, percentage Ramachandran outliers, and percentage of bad side-chain rotamers. Removing MolProbity score and Ramachandran favored residues reduced the collinearity of the other variables to moderate values (1.0568 to 3.3434). Two principal components were sufficient to explain 72.18% of the variation between the models’ metrics (**Figure 6**).

The results of the PCA indicate that the analyses captured the multi-dimensional variation in the metrics, and the presence of broad, systematic differences between how the methods handle stereochemistry which may affect the biological relevance of the generated structures. Firstly, KMeans clustering (*k* = 5) revealed a high degree of separation by model with distinct separation between the structures generated by ESMFold, Boltz, and the AlphaFold methods, but little to no separation between AlphaFold 2 and AlphaFold 3. The coefficients of the principal components also suggest that the percentage of Ramachadran outliers (coef. = 0.6333), and rotamer outliers (coef. = 0.6555) were driving the variance in the first component whereas the second principal component was largely driven by the clash score (coef. = 0.8415), and the third was most influenced by the Cβ deviations (coef. = 0.78922).

The correlation between the structures’ confidence scores and the steric features was also investigated to determine if confidence is indicative of major steric hindrances. Ideally, the methods would have less confidence (lower pLDDT scores) for structures that are physically or chemically implausible. This inverse relationship trend was seen in the data (**Figure 7**), particularly in the MolProbity Scores. Notably, while the relationship between the MolProbity and pLDDT scores was linear for the MSA-based methods (AlphaFold 2, AlphaFold 3, Boltz-1, and Boltz-2), this relationship was polynomial for ESMFold. This suggests the presence of another limiting factor that impacts this relationship, such as the representation of similar sequences in the training data.

**Figure 7.**
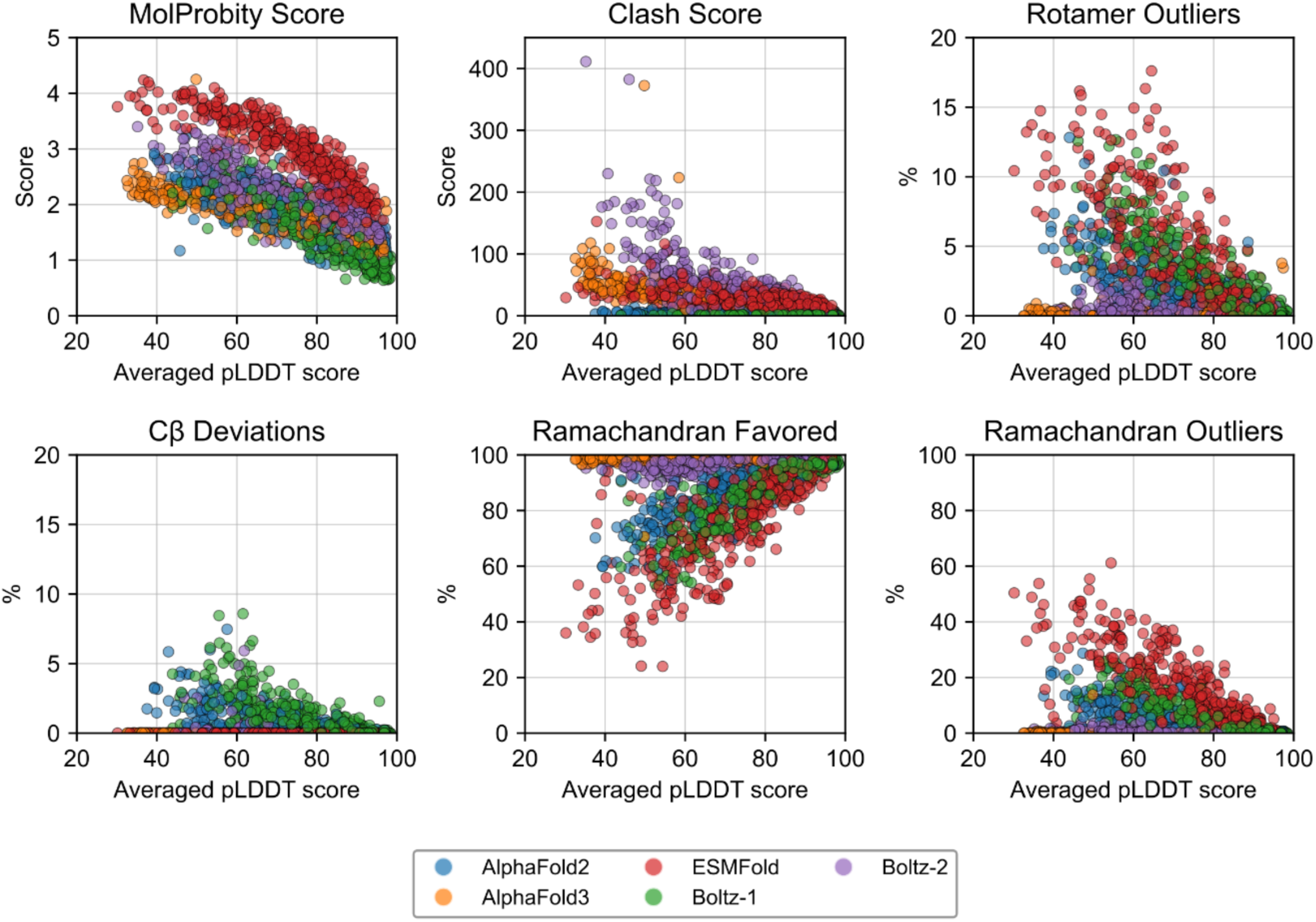
The relationship between the pLDDT score and steric features of the generated structures, by structure prediction program.

### A case study using experimentally validated structures from the PDB

Global and local structural alignments were performed to assess the alignment of the generated structures with experimentally validated structures from the PDB (**Figure 8**). The program Foldseek was selected to align structures primarily due to its speed and diverse outputs/output parameters. There are very few experimentally confirmed protein structures for maize in the PDB (272 structures as of April 2025) and just eight experimentally confirmed structures amongst our benchmark classical gene set. Thus the analysis was expanded to comparisons of the experimental and computational structures from *Arabidopsis thaliana* (Arabidopsis), *Oryza sativa* (rice), *Triticum aestivum* (bread wheat), and *Glycine max* (soybean).

**Figure 8.**
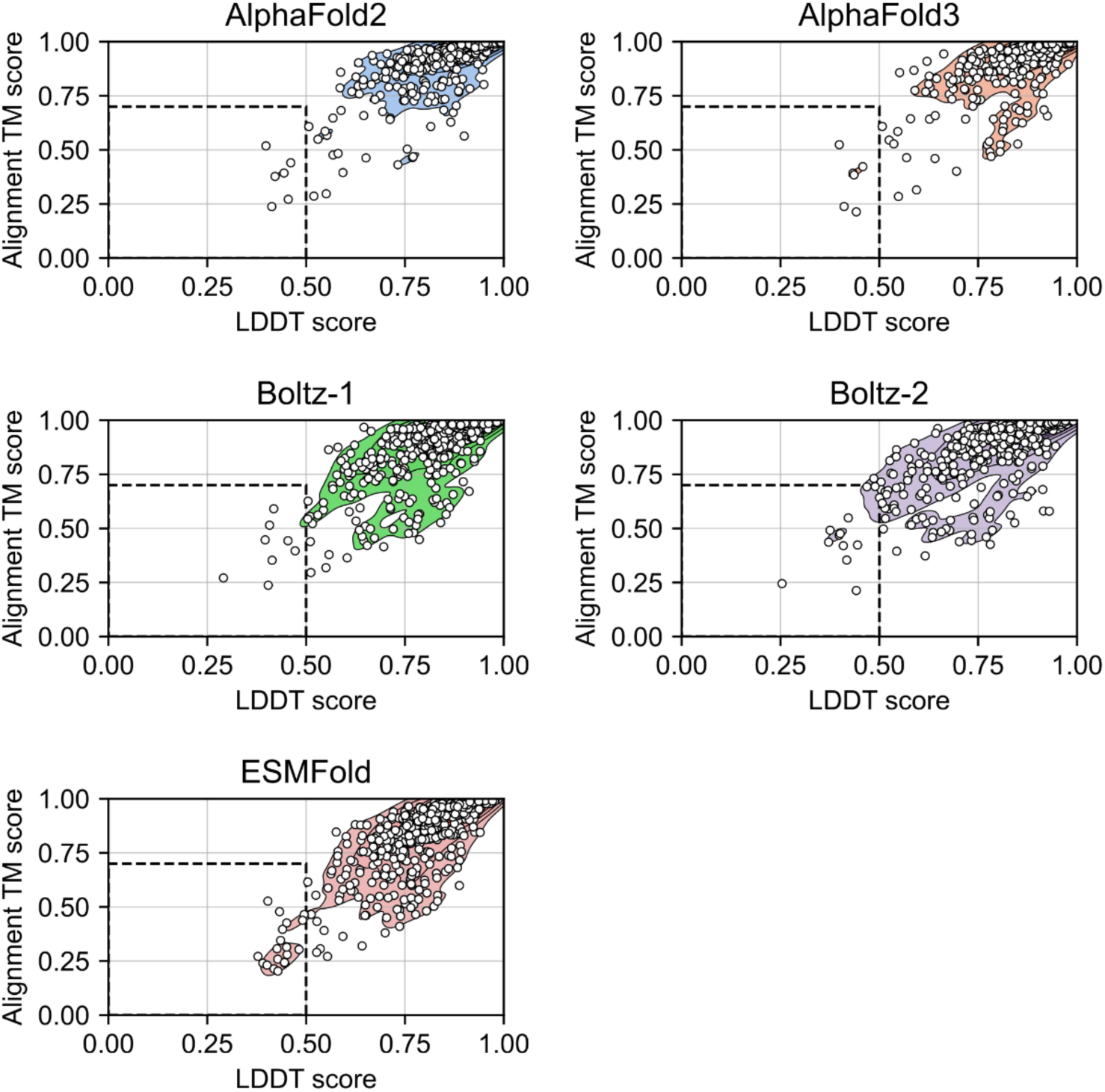
An evaluation of experimental and *in silico* plant protein structures generated by AlphaFold 2, AlphaFold 3, Boltz-1, Boltz-2, and ESMFold. Distributions of global and local structural alignments (**left**) are shown for experimentally validated plant proteins from the Protein Data Bank (PDB) and corresponding *in silico* structures predicted by the four folding methods. In each panel, the x-axis represents the local structural alignment as the lDDT score whereas the y-axis depicts the global structural alignment via the TM score. Overall, the models performed well, generating structures with high local (LDDT ≥ 0.7) and global (TM-score ≥ 0.5) similarity to their experimental counterparts. Structures with poor alignment (in the dashed sections) were further analyzed using the SWISS-MODEL Structure Assessment tool to examine steric features.

There was a high degree of global (TM score ≥ 0.50) and local (LDDT score ≥ 0.70) structural agreement for most of the methods. Overall, 15.7% (76/485) of the bread wheat structural comparisons indicated either poor global or local agreement, compared to 9.2% (255/2,775) from Arabidopsis, 6.7% (28/421) from maize, 4.2% (7/167) from soybean, and 2.2% (6/269) in rice.

Initially, the distributions between each plant species also did not seem to significantly differ, though Arabidopsis, bread wheat, and maize seemed to have more discrepancies in the structural alignments than rice or soybean (**Supplemental Figure 2**). To evaluate whether structural prediction programs differed in their performance across species, we analyzed three structural quality metrics – LDDT, Alignment TM Score, and RMSD – using linear mixed effects models. Specifically, we sought to test the hypothesis that the models performed equally well in assembling the structure of plant proteins in the PDB. We selected these metrics because of their prominence in the literature where they often serve as indicators to evaluate local structural alignments (LDDT) and global structural alignments (TM Score, RMSD) between proteins. Thus, for each species and alignment metric, we fit separate linear mixed models with the program as a fixed effect and protein as a random effect (to control for proteins that are intrinsically harder to predict). Where appropriate, pairwise post-hoc comparisons between programs were conducted using Welch’s t-tests to account for unequal variances between groups. To control for multiple comparisons, Bonferroni correction was applied to the resulting *p*-values. Comparisons were considered statistically significant at an adjusted p < 0.05.

The resulting p-values for the pairwise comparisons of the programs’ R.M.S.D. by species are shown in **Figure 9**; comparisons were also calculated between the models LDDT scores (**Supplemental Figure 3**) and TM scores (**Supplemental Figure 4**). AlphaFold 2 and AlphaFold 3 tended to outperform the other models for Arabidopsis and wheat proteins, but there were no statistically significant differences between the R.M.S.D. for most of the programs for maize, rice, or soybean proteins. This trend was similar for the TM scores, but there were more statistical differences between the models and species for the alignments’ LDDT scores. At present, however, there isn’t enough information to further discern the reason for these discrepancies in local structure between the computational and experimental structures. For a more comprehensive analysis, more structures are needed.

**Figure 9.**
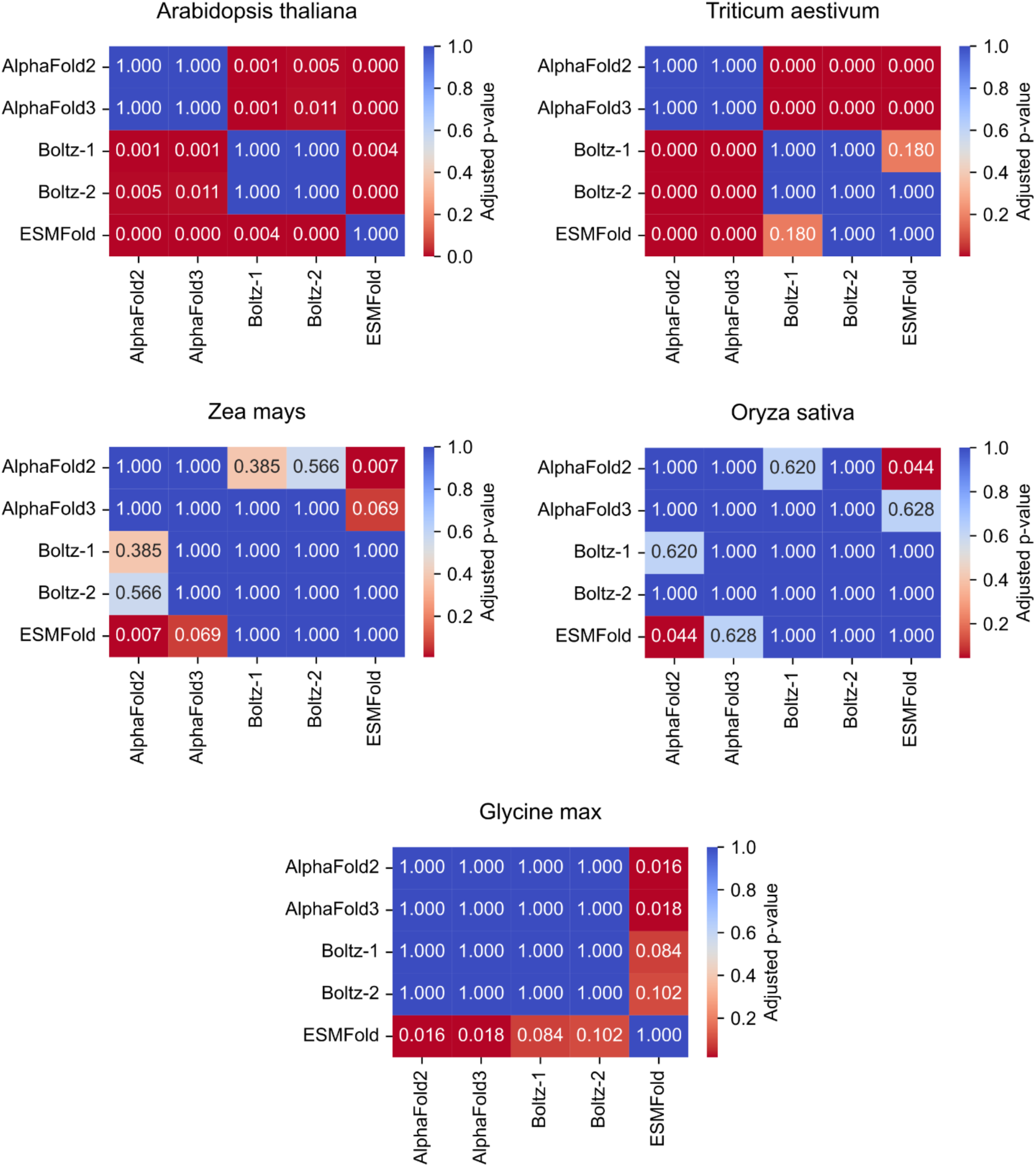
Heatmaps showing pairwise adjusted p-values from post-hoc pairwise t-tests comparing the **RMSD** between experimental and computational structures generated for each prediction program, by species. Each panel represents one species, with cells annotated by Bonferroni-adjusted p-values for all program pairs. Lower p-values (darker colors) indicate statistically significant differences between programs. This visualization highlights which program pairs differ significantly in their performance within each species and metric.

We identified 86 *in silico* structures that exhibited poor global and local alignments with their experimental counterparts. These structures were further analyzed for steric hindrances, with the hypothesis that these poor alignments may be due to more severe steric clashes. Notably, there were only seven gene models for which all four methods generated computational structures with poor global and local alignment to the ground truth PDB structure: maize (Zm00001d019033), wheat (TraesARI6D03G03676800, Traes5BL3608AAB59, TraesARI4A03G02187480), and Arabidopsis (At3g16450, At5g43410, At3g55470). However, by superimposing the values of the steric features onto the prior distributions from **Figure 6**, the poor alignments do not appear to have a higher prevalence of steric hindrances than the proteins from the classical gene models (**Figure 10**, **Supplemental Figure 3**). Overlaying the new coordinates in the principal component analysis also shows how the computational structures, despite poor alignment with the experimental structures, still align well with what is expected regarding their steric features (**Figure 10)**. Interestingly, however, a few of the experimental structures did not fit within the distribution, as evidenced by the PCA (**Figure 10**). Further experimentation to clarify this discrepancy would be difficult, however, due to the low number of experimentally resolved plant protein structures in the PDB.

**Figure 10.**
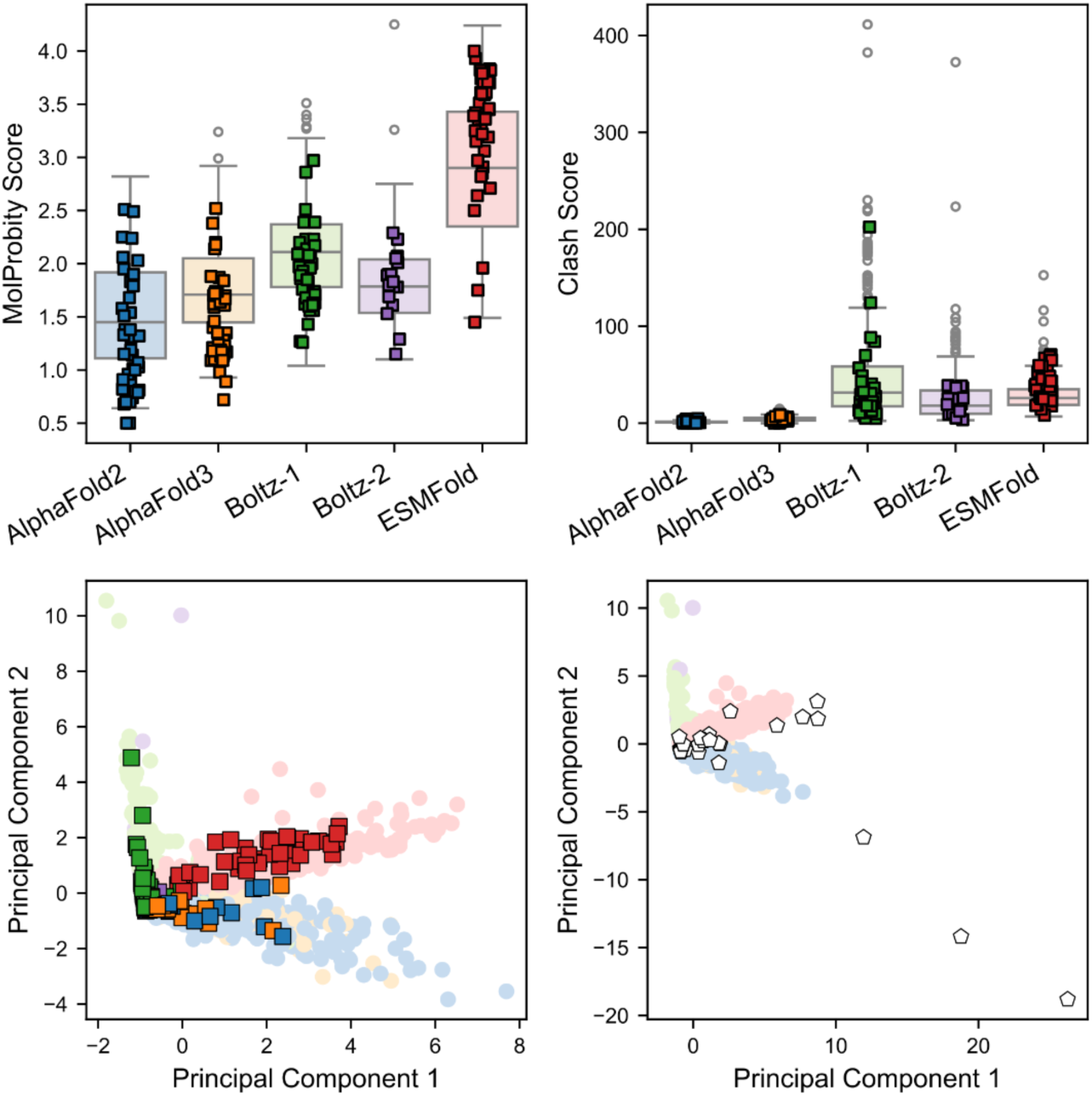
The distribution of steric features and principal component analysis (PCA) of structures from classical maize genes, with the positions of poorly aligned predictions superimposed. The top panels depict the distribution of the MolProbity score and Clash score of the classical genes in lighter colors, and the distribution of the metrics for the poorly aligned structures in darker colors. Although these outliers were inaccurate in terms of structural alignment, their steric features generally remained within the expected distribution (**top**). The lower panels show the PCA from the classical maize genes overlapped by the observations from the poor alignments (**bottom left)** or the ground truth experimental protein structures (**bottom right**). Several observations from the experimental structures fall outside the original PCA distribution **(bottom right**).

## CONCLUSION

Maize is one of the world’s most important agronomic crops and has extensive, publicly available genomic resources. However, it remains severely underrepresented in protein structure databases that are used to train many structure prediction tools. This benchmarking study offers a comprehensive evaluation of five state-of-the-art protein structure prediction tools (AlphaFold 2, AlphaFold 3, ESMFold, Boltz-1, and Boltz-2) within the context of maize proteomics. These models represent significant advances in structural biology that could supplement current protein engineering initiatives to improve grain crop pathogen resistance^44,45^, abiotic tolerance^46^, herbicide tolerance^47^, and enhance nutritional quality^46,48^.

Our findings reveal limitations in their application to plant-specific proteins, particularly those that are evolutionarily young or species-specific. Our phylostratigraphic approach revealed a clear trend: proteins encoded by evolutionarily young gene models that were specific to maize or the Poaceae family were predicted with lower structural confidence. This pattern was robust across both AlphaFold 2 and ESMFold models, and was further supported by Kruskal-Wallis non-parametric analyses and post-hoc pairwise comparisons. Importantly, the decline in model confidence for young genes may reflect the structural novelty and rapid evolution of these proteins, which are often involved in adaptive processes such as pathogen response and abiotic stress tolerance. Thus, despite their agronomic importance, these proteins remain structurally under-characterized due to their poor representation in training data and limited homology to resolved structures.

The benchmarking study also emphasized the importance of resolving and functionally characterizing proteins with limited sequence or structural conservation. Proteins annotated with both Pfam and CATH domains, which we used as indicators of sequence and structure conservation, respectively, consistently yielded the most confident predictions. By contrast, proteins lacking either domain exhibited substantially lower confidence, with the absence of both domains compounding this effect. These results highlight the current reliance of structure prediction algorithms on well-characterized proteins with conserved features, which are more prevalent in bacterial, fungal, and vertebrate datasets than in plant datasets.

The stereochemical assessments, particularly the distinct clustering patterns in the Principal Component Analysis (PCA), suggest that each method imparts a characteristic steric “signature” onto its predictions, which could be used to inform quality control protocols or model selection in future applications. According to the linear mixed-effects modeling, AlphaFold 2, AlphaFold 3, and Boltz-2 produced the most stereochemically sound structures overall. AlphaFold 3 achieved lower frequencies of rotamer outliers and Cβ deviations than AlphaFold 2, suggesting modest gains in physical plausibility despite slightly reduced pLDDT scores. Similarly, Boltz-2 structures experienced many fewer occurrences of clashing compared to Boltz-1 structures without significant increases in structure generation time. ESMFold and Boltz-1, while advantageous in terms of speed, frequently generated structures with elevated clash scores and Ramachandran outliers, highlighting their trade-off between computational efficiency and physical accuracy. Boltz-2 provides a good middle ground in terms of both speed and confident predictions, with few stereochemically sound structures overall. AlphaFold 2 remains the overall best method in terms of predicting the most accurate protein structures, but comes at a steep cost in terms of needing large GPU resources for most proteomes.

Taken together, our results highlight the substantial challenges that remain when applying general-purpose protein structure prediction tools to evolutionarily novel or taxonomically restricted plant genes. The dual limitations of confidence and stereochemical quality for these proteins suggest that structure prediction models are not yet fully generalizable across the diversity of plant proteomes. This has significant implications for fields such as crop improvement and synthetic biology, where structurally modeling lineage-specific proteins could drive innovation but is currently hindered by inadequate tool performance.

To address these limitations, we recommend a multi-pronged approach: (1) experimental efforts to resolve the structures of orphan, young, and lineage-specific plant proteins, particularly those implicated in stress adaptation; (2) increased representation of plant proteins in structural databases like the Protein Data Bank; and (3) incorporation of diverse plant-specific training data into future versions of structure prediction models. The development of plant-specific benchmarking datasets and quality control standards will also be critical to guide model refinement and validation. Ultimately, bridging the gap between structure prediction capabilities and the biological complexity of plant proteomes is essential for realizing the full potential of computational structural biology in agriculture.

## Data availability

The generated structures are available at MaizeGDB (maizegdb.org), and the code used in this study is available at GitHub https://github.com/Maize-Genetics-and-Genomics-Database/Structural-Assessment-Maize-CaseStudy

## Acknowledgements

The work for this project was performed on the Atlas and Ceres high-performance clusters as part of the USDA-ARS SCINet initiative. We would like to thank the SCINet administrative staff and the Virtual Research Support Core team. This work was supported by the U.S. Department of Agriculture, Agricultural Research Service. Mention of trade names or commercial products in this publication is solely for the purpose of providing specific information and does not imply recommendation or endorsement by the U.S. Department of Agriculture. USDA is an equal opportunity provider and employer. We would also like to thank Austin Weigel for his guidance in performing the structural assessments.

## Supplemental Tables and Figures

**Supplemental Table 1.**
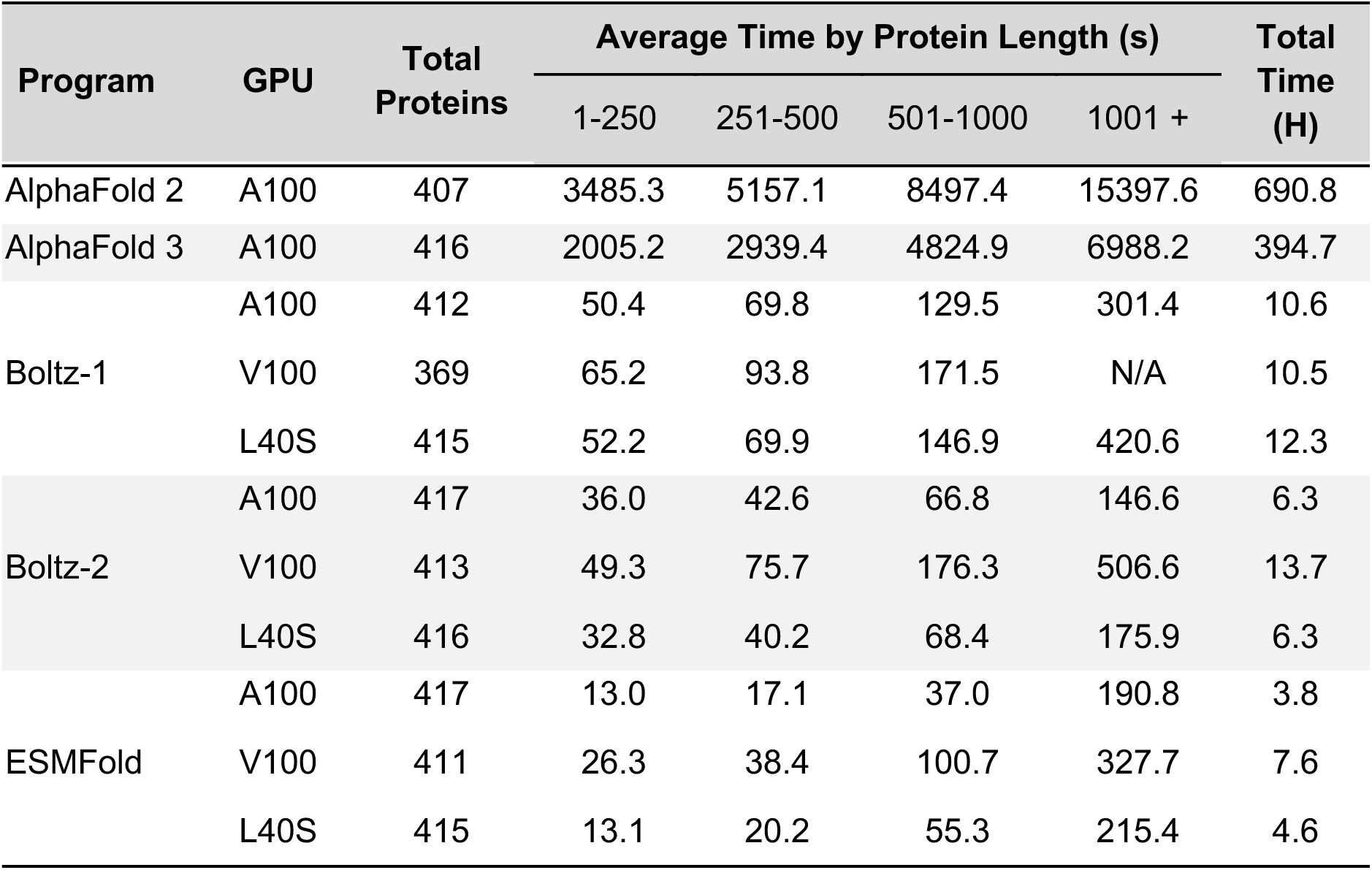
A tabulated summary of the runtimes for each model on A100, V100, and LS40 GPU cores by protein length.

**Supplemental Figure 1.**
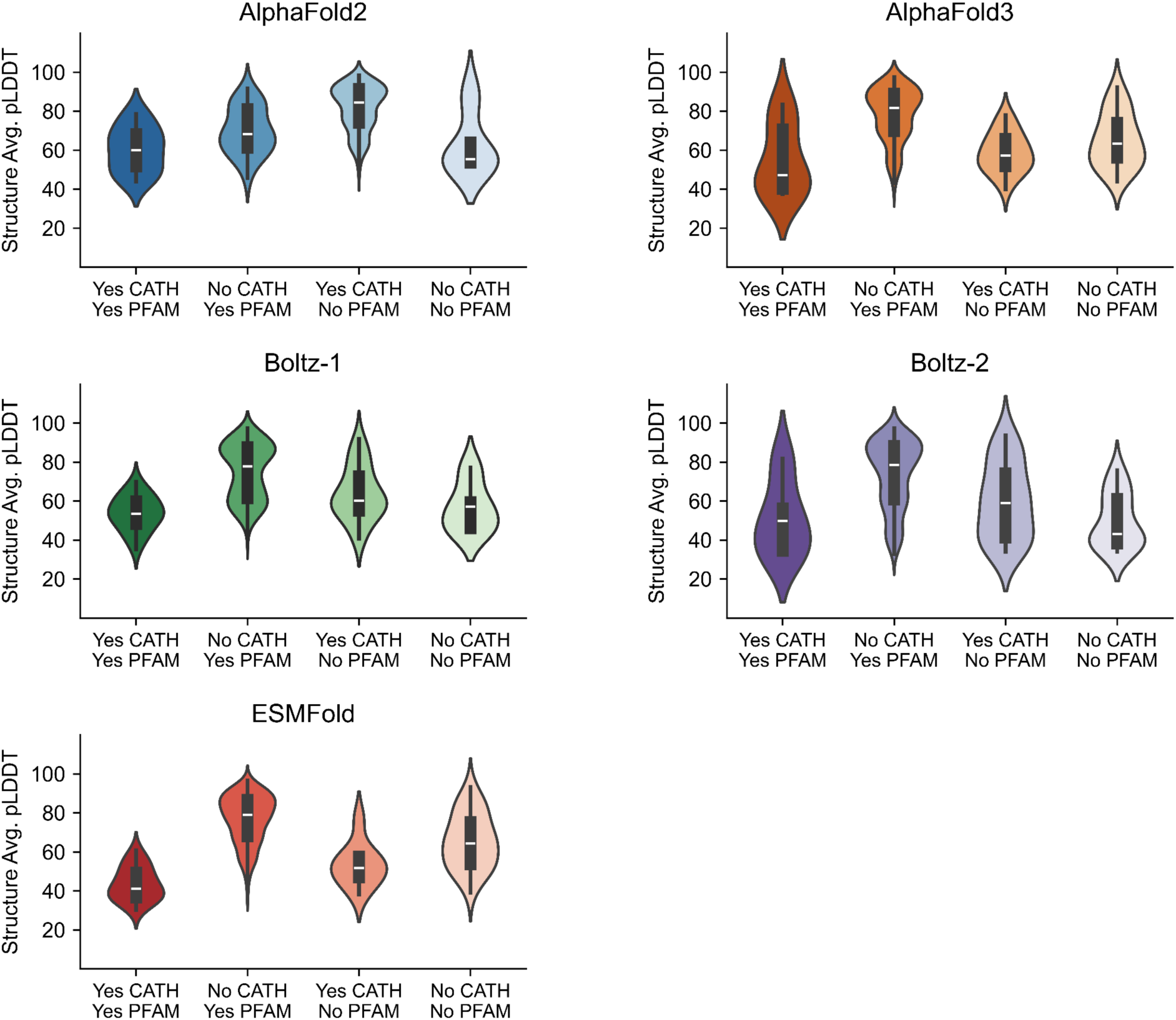
A comparison of the structure confidence scores, by absence/presence of conserved sequence and structural domains.

**Supplemental Figure 2.**
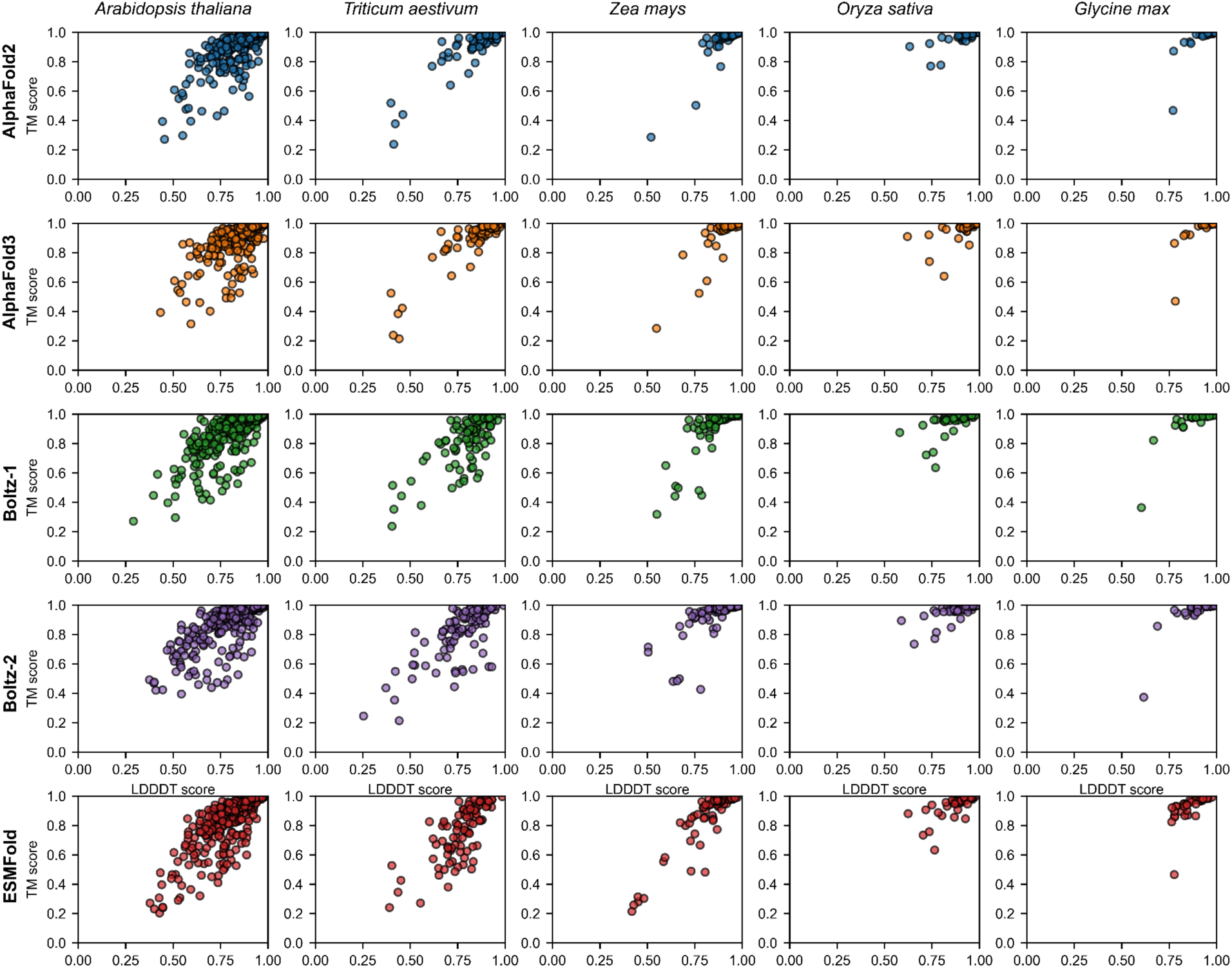
Global and local structural alignments between experimental and *in silico* plant protein structures generated by AlphaFold 2, AlphaFold 3, Boltz-1, and ESMFold, by species.

**Supplemental Figure 3.**
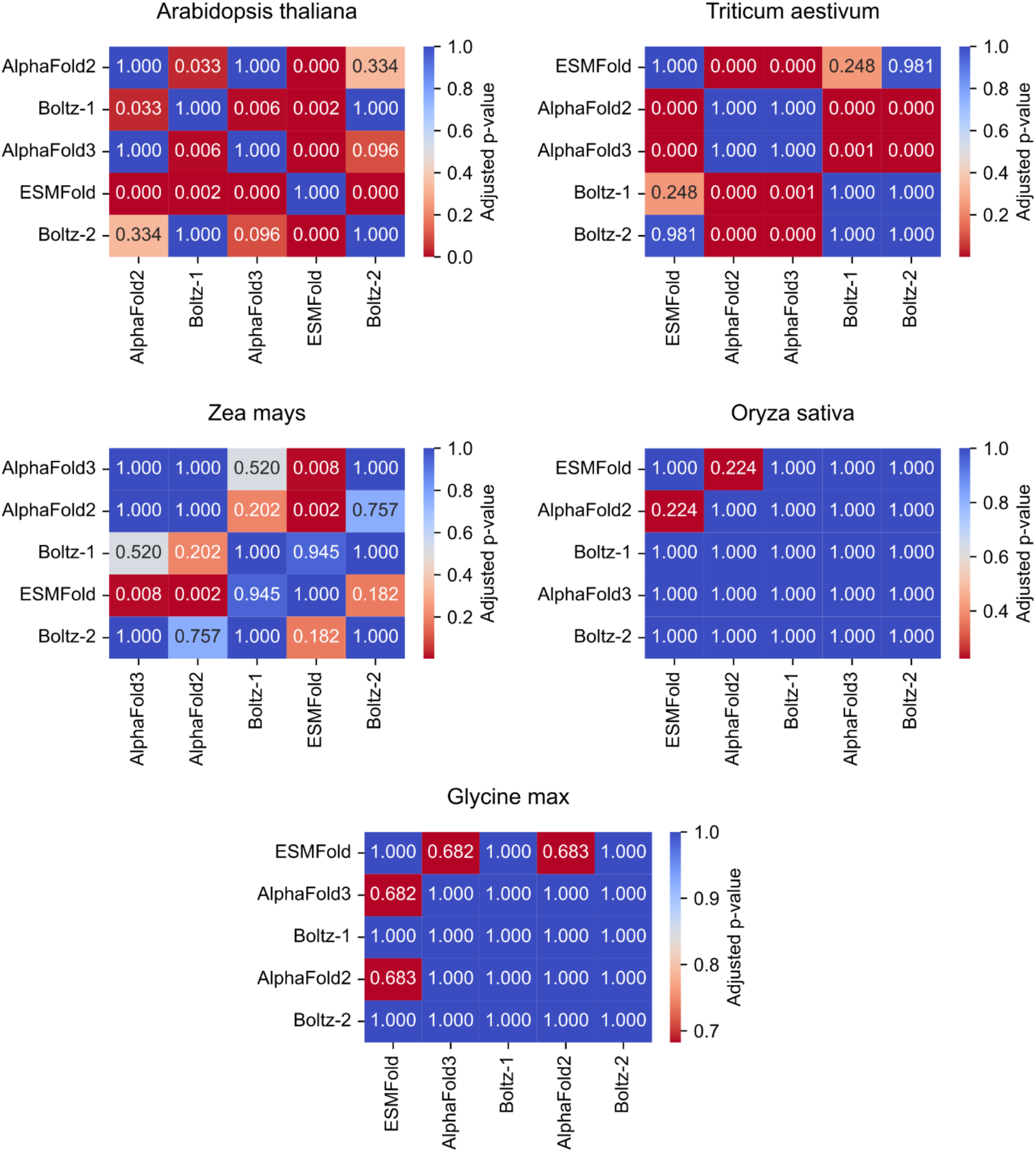
Heatmaps showing pairwise adjusted p-values from post-hoc pairwise t-tests comparing the **alignments’ TM score** between experimental and computational structures generated for each prediction program, by species. Each panel represents one species, with cells annotated by Bonferroni-adjusted p-values for all program pairs. Lower p-values (darker colors) indicate statistically significant differences between programs. This visualization highlights which program pairs differ significantly in their performance within each species and metric.

**Supplemental Figure 4.**
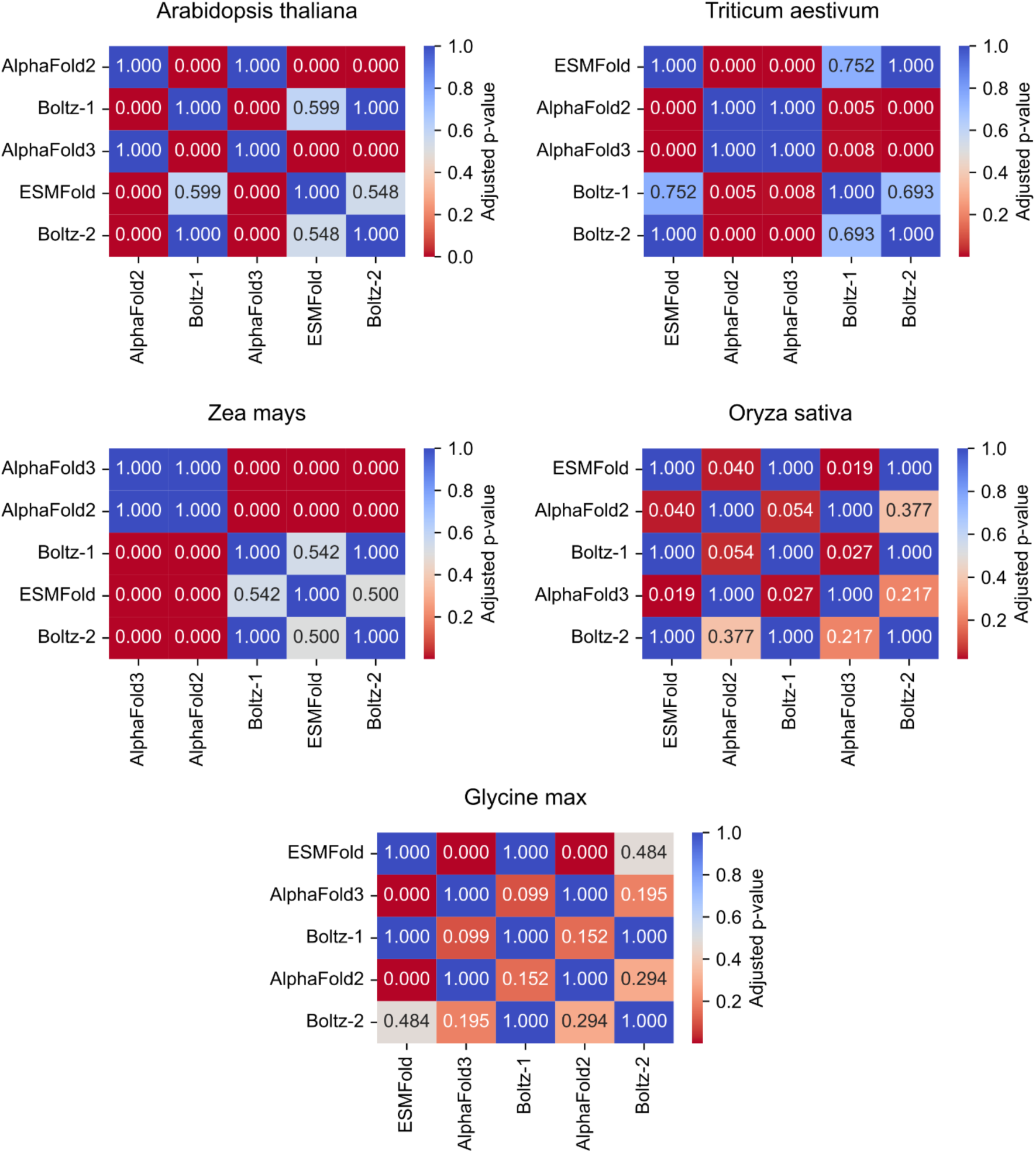
Heatmaps showing pairwise adjusted p-values from post-hoc pairwise t-tests comparing the **alignments’ LDDT score** between experimental and computational structures generated for each prediction program, by species. Each panel represents one species, with cells annotated by Bonferroni-adjusted p-values for all program pairs. Lower p-values (darker colors) indicate statistically significant differences between programs. This visualization highlights which program pairs differ significantly in their performance within each species and metric.

**Supplemental Figure 5.**
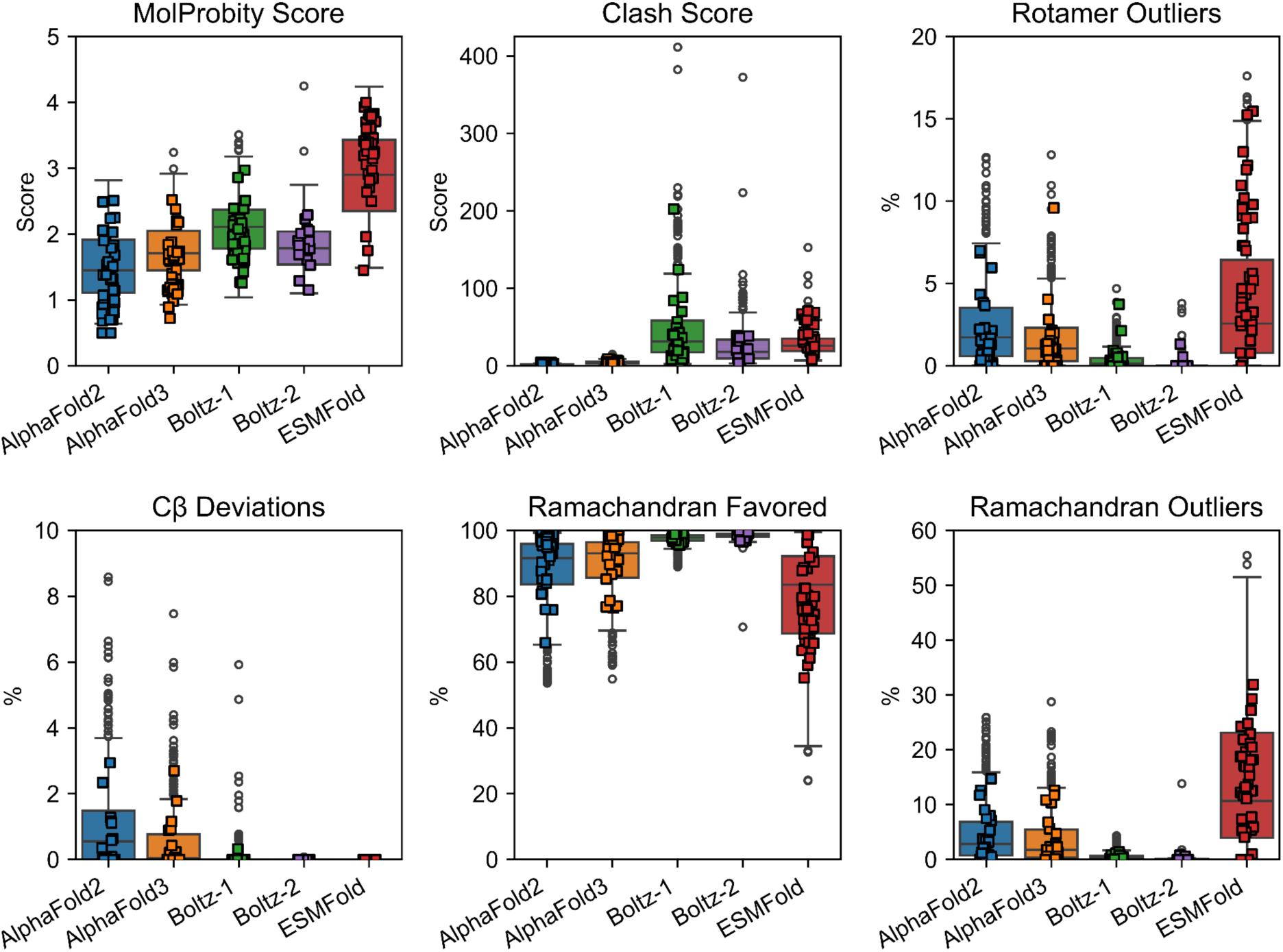
Structural assessments of the plant PDB proteins folded using AlphaFold 2, AlphaFold 3, Boltz-1, Boltz-2, or ESMFold.

